# The Effects of RNA.DNA-DNA Triple Helices on Nucleosome Structures and Dynamics

**DOI:** 10.1101/2022.07.02.498574

**Authors:** Havva Kohestani, Jeff Wereszczynski

**Affiliations:** Illinois Institute of Technology, Department of Biology; Illinois Institute of Technology, Departments of Physics & Biology

## Abstract

Non-coding RNAs (ncRNAs) are an emerging epigenetic factor and have been recognized as playing a key role in many gene expression pathways. Structurally, binding of ncRNAs to isolated DNA is strongly dependent on sequence complementary, and results in the formation of an RNA.DNA-DNA (RDD) triple helix. However, in vivo DNA is not isolated, but is packed in chromatin fibers, the fundamental unit of which is the nucleosome. Biochemical experiments have shown that ncRNA binding to nucleosomal DNA is elevated at DNA entry and exit sites and is dependent on the presence of the H3 N-terminal tails. However, the structural and dynamical bases for these mechanisms remains unknown. Here, we have examined the mechanisms and effects of RDD formation in the context of the nucleosome using a series of all-atom molecular dynamics simulations. Results highlight the importance of DNA sequence on complex stability, elucidate the effects of the H3 tails on RDD structures, show how RDD formation impacts the structure and dynamics of the H3 tails, and show how RNA alters the local and global DNA double helical structure. Together, our results suggest ncRNAs can modify nucleosome, and potentially higher-order chromatin, structures and dynamics as a means of exerting epigenetic control.

**SIGNIFICANCE:** Non-coding RNAs (ncRNAs) play an essential role in gene regulation by binding to DNA and forming RNA.DNA-DNA (RDD) triple helices. In the cell, this occurs in the context where DNA is not a free double helix but is instead condensed into chromatin fibers. At the fundamental level, this compaction involves wrapping approximately 147 DNA basepairs around eight histone proteins to form the nucleosome. Here, we have used molecular dynamics simulations to understand the interplay between the structure and dynamics of RDD triple helices with the nucleosome. Results highlight the importance of RNA sequence on RDD stability regardless of its environment and suggest potential mechanisms for cross-talk between epigenetic factors and the effects of ncRNA binding on local and higher-order chromatin structures.

## 1 INTRODUCTION

Epigenetic regulation mechanisms are critical in inducing heritable phenotype changes in biological systems without altering their core genetic DNA sequences. In vivo, reversible epigenetic mechanisms engage various molecular structures from small post-translational modifications (PTMs) to RNAs to large protein complexes. For example, chromosome X-inactivation, histone deacetylation, histone modification, DNA methylation, and noncoding RNA-mediated (ncRNA) processes are a few examples of epigenetic control mechanisms (1–5). In particular, experiments have shown the involvement of different ncRNAs like Piwi-interacting RNAs (piRNAs), microRNAs (miRNA), and small interfering RNAs (siRNAs) in embryonic development and diseases such as cancer. For example, there is an association between active mRNA transcription sites and trimethylation of lysine 4 (H3K4me3) and lysine 36 of histone H3 (H3K36me3). In contrast, monomethylation of lysine 4 (H3K4m1) and acetylation at lysine 27 in H3 (H3K27ac) is a sign of transcription upregulation from enhancer RNA (eRNA) activated promoters (6, 7). These factors suggest RNA-based molecular markers may be useful in the design of molecular markers for the detection of targeting cancer cells and that ncRNAs may have therapeutic applications in eukaryotic systems by inducing viral resistance (5–10).

Structurally, noncoding RNAs fit into two categories based on nucleotide size. Long noncoding RNAs (lncRNAs) are composed of a minimum of 200 nucleotides, while RNAs with 20-30 nucleotides are considered small ncRNAs (8, 11). The activity and influence of small or long ncRNAs depends on several factors such as their localization, and the type of RNA-DNA or RNA-protein interactions they make (9, 11–13). Various elements play a role in initiating a successful interaction between ncRNAs and genomic DNA to create functional RNA.DNA-DNA (RDD) triple helices. Experimental evidence has shown that the formation of proper hydrogen bonding between triplex-forming oligonucleotides (TFOs) and a DNA double helix are highly sequence-dependent where these factors have a preference to bind to DNA at specific triplex targeting sites (TTS) that have a higher presence at an enhancer or promoter locus (13–19). Besides the significant sequence specificity, efficient binding of RNA to a DNA counterpart in solution is reliant on factors such as the RNA length, C-G content of the TTS, the parallel or anti-parallel alignment of nucleotide strands, and the presence or absence of point mutations (17–21).

In vivo, ncRNAs exert their influence by interacting with DNA that is packaged into chromatin fibers. The fundamental building block of chromatin is the nucleosome, a complex consisting of two copies of the H4, H3, H2A, and H2B histones, along with approximately 145-147 DNA basepairs (22). Structurally, each core histones has an unstructured N-terminal tail of varying lengths, three a-helices which are connected by two loops, and H2B also has a short and unstructured C-terminal tail (23). The histones assemble into a disc-like structure around which the DNA wraps (Figure 1) (24, 25). The nucleosome itself is a highly dynamic entity, and adopts an ensemble of structures characterized by core histone, histone tail, and DNA breathing motions (24–28). Other epigentic factors, such as reader domain binding, linker histone interactions, and the replacement of canonical histones with variants have been shown to influence these dynamics as a way to influence gene expression and higher-order chromatin structures (28–32).

**Figure 1:**
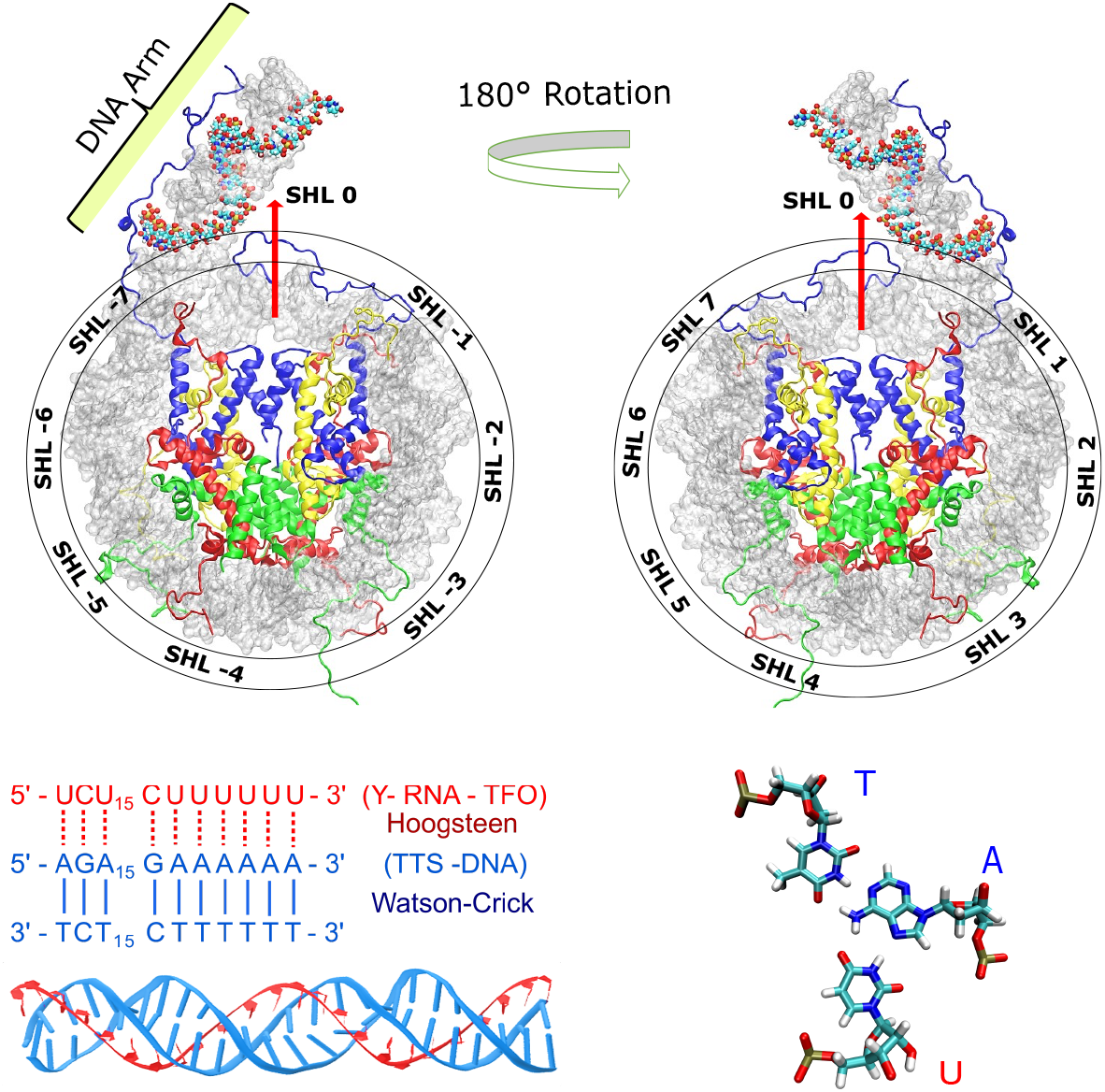
(Top) A general view of the nucleosome and its structure. The H3 (blue), H4 (yellow), H2A (red) and H2B (green) histones form the protein core around which approximately 147 basepairs of DNA (silver) wrap. Here, there are an additional 20 DNA basepairs at the entry DNA site, which we term the DNA arm. The reference dyad axis is shown by a red arrow. By convention, DNA locations are denoted by their superhelical locations (SHL) relative to the dyad, with DNA closer to the entry site having SHLs less than 0 and closer to the exit site greater than 0. (bottom left) Pyrimidine RNA triplex forming oligonucleotides have a preference for A-T rich TTS. A RNA-TFO is represented in red against a blue TTS-DNA sequence (17). (bottom right) Hoogsteen base-pairs help to stabilize RNA.DNA-DNA triple helices and we have selected a parallel (Y) RNA-TFO sequence to maximize the complementary binding between the RNA and DNA double helix.

To understand how ncRNAs interact with DNA in chromatin, Maldonado *et al*. recently examined the stability of RDD triple helices in nucleosome structures. In particular, electrophoretic mobility shift assays showed that formation of a stable RDD triple helix occurs primarily at the entry/exit sites of the nucleosomal DNA. In addition, the H3 tails were shown to play a key role in increasing RDD stability in comparison to truncated nucleosome which did not include these long, disordered regions (17, 33). However, the molecular processes underlying these mechanisms remains elusive, as there are no published crystal structure available for RDD helices in chromatin structures, and there is little information about how RNA binding affects nucleosome dynamics.

Here, we have sought to examine the molecular mechanisms of RDD triple helices when bound to nucleosome systems. To do so, we generated models of nucleosome structures with an extended DNA arm at the entry DNA site to which a TFO RNA could bind. We performed a series of molecular dynamics (MD) simulations of isolated RDD triplexes and full-length and H3-tailless nucleosomes containing RDD triplexes with either TTS or NoTTS DNA sequences in the DNA arm. Results are in agreement with experiments which have shown that the stable formation of an RDD triplex is driven by the DNA sequence, as in all cases of NoTTS containing models there was unstable RNA binding regardless of its environment. Furthermore, the H3 tails are shown to affect RNA binding and adopt distinct conformations in the presence of the RDD triplexes. Finally, analyses of the DNA nucleic acid parameters show that RNA binding induces a transition to an intermediate structure between B-and A-form. Overall, these results demonstrate that RDD triplex formation is largely dependent on the correct positioning of the TTS sequence and suggest that RDD binding may influence other epigentic factors, particularly those involving the H3 tails.

## 2 MATERIALS AND METHODS

### 2.1 System Setup

We designed five nucleosome systems based on the protein core of the 1KX5 and DNA of the 6VYP protein data bank crystal structures (Table S1) (23, 34). 6VYP was stripped of all protein segments. We kept only one DNA arm to avoid potential interactions of the two extended DNA arms in the simulation box. The sequence of the entry DNA arm was mutated in the w3DNA webserver to match the En3-TTS sequence for the samples with the TTS sequences (17, 35). Experimental studies have suggested that an RNA-TFO should be a minimum of 19 nucleotides in length to generate an RDD triplex, while increases in the RNA length beyond 27 bases do not enhance its DNA affinity (18). As a middle ground, we constructed a 24 basepair RNA-TFO in the 3D-NuS webserver for the triplex targeting and non-targeting sequences (36). RDD triplexes were replaced and adjusted at the DNA arm via Chimera’s modeller function (37). The canonical protein core was added to the final mutated edited nucleic acid structure and assembled in Visual Molecular Dynamics (VMD) and Chimera (37, 38). The model pool consisted of seven systems, including isolated TTS and NoTTS triplexes, TTS-nucleosomes with and without H3 tails, NoTTS-nucleosomes with and without H3 tails, and a nucleosome model without RNA. For clarity, we have used superscript ^*tH*3^ for nucleosomes with truncated H3 tails and labeled DNA locations by their Superhelical location (SHL), which range from -7 to 7 with a value of SHL=0 corresponding to the nucleosome dyad (Figure 1) (25, 39–41).

All systems were built in the tleap tool of the Amber molecular dynamics package (42, 43). The Amber ff14SB, OL15, OL3, OPC force fields were used for the protein, DNA, RNA, and water force fields, while ions parameters were based on the Joung and Cheatham model (44–49). All systems were neutralized and solvated in a water box with a 12 Å solvent buffer and an additional 0.15 M NaCl concentration. The ParmEd (parameter file editor) program was used to repartition the hydrogen atom masses to reduce high-frequency motions, which allowed the use of a 4ps MD timestep instead of 2ps (50). All systems were minimized twice for 10000 steps, with and without heavy atom restraints. Systems were then gradually heated in the canonical ensemble from 0 K to 300 K over 5000 steps with restraints on the solute heavy atoms. Restraints were released by relaxing the systems in the isothermal-isobaric ensemble over 300 ps. The Langevin piston was used for a barostat with Langevin dynamics for temperature control with a collision frequency of 2 ps^−1^ (51). All simulations were performed with the GPU accelerated version of PMEMD for three replicas of each system (52). Table S1 represents a summary of the simulations performed.

### 2.2 Simulation Analyses

CPPTRAJ was used to measure root-mean-square-deviations (RMSDs), root-mean-square-fluctuations (RMSFs), inter and intramolecular distances, and various nucleic acid parameters (53). For all analyses, 100 ns of simulation time was taken for equilibration. VMD was used to calculate the number of hydrogen bonds and also produce figures (38), and python was used for data processing and generating plots (54). The MM/GBSA approach was used to estimate the interaction energy between the H3 histones and nucleic acid helices (55, 56). The Generalized Born model with *igb*=5 implicit solvent models was utilized for the polar term along with a salt concentration of 0.15 M and the *mbondi2* radii set (57, 58). For interpretation, the energy terms were grouped into two terms: Δ*E*_*vd*_*w* and Δ*E*_*vd*_*w*. Δ*E*_*vd*_*w*, is the sum of the molecular mechanics van der Waals energy and the solvation apolar energy differences, and *E*_*elec*_ is the sum of the molecular mechanics electrostatics and polar solvation energy differences. A residue decomposition of the MM/GBSA analysis was performed to estimate the energy values for each protein residue interacting with the RDD.

## 3 RESULTS

### 3.1 Triplex targeting sites in nucleosomes form stable sequence dependent RDD helices

Seven sets of simulations were performed, which included isolated TTS-DNA and noTTS-DNA RDD systems, nucleosome/RNA systems with and without the TTS-DNA and with and without the H3 tails, and a canonical nucleosome without RNA (Table S1). For each system, five 1.1 µs simulations were performed. Simulations showed that a TTS-DNA sequence is required for maintaining the stability of an RDD triplex, regardless of its environment. In simulations of the isolated triplex with the TTS sequence, the nucleic acid root-mean-square-deviation (RMSD) remained relatively low and constant, with values of ∼2-4 Å (Figure 2a). In contrast, for isolated triplexes with a non-TTS DNA sequence the system quickly destabilized, typically in the first 200 ns, and reached RMSD values in the range of 40 Å (Figure 2b). In the context of the full-length nucleosome, the TTS triplex stayed well formed with RMSD values of 2-6 Å, whereas some destabilization of the triplex was noted in the non-TTS triplex, with one simulation sampling RMSD values of ∼20 Å (Figure 2d). Removal of the H3 tails resulted in similar sampling of the TTS triplex (Figure 2e), however there was a marked increase in the RMSD of the non-TTS sequence to values as large as 40 Å (Figure 2f). In each of these systems the RNA component of the RDD triplex was the most dynamic, with trends in RNA RMSD values largely mirroring the overall trends in the RDD RMSD values (Figure S1).

**Figure 2:**
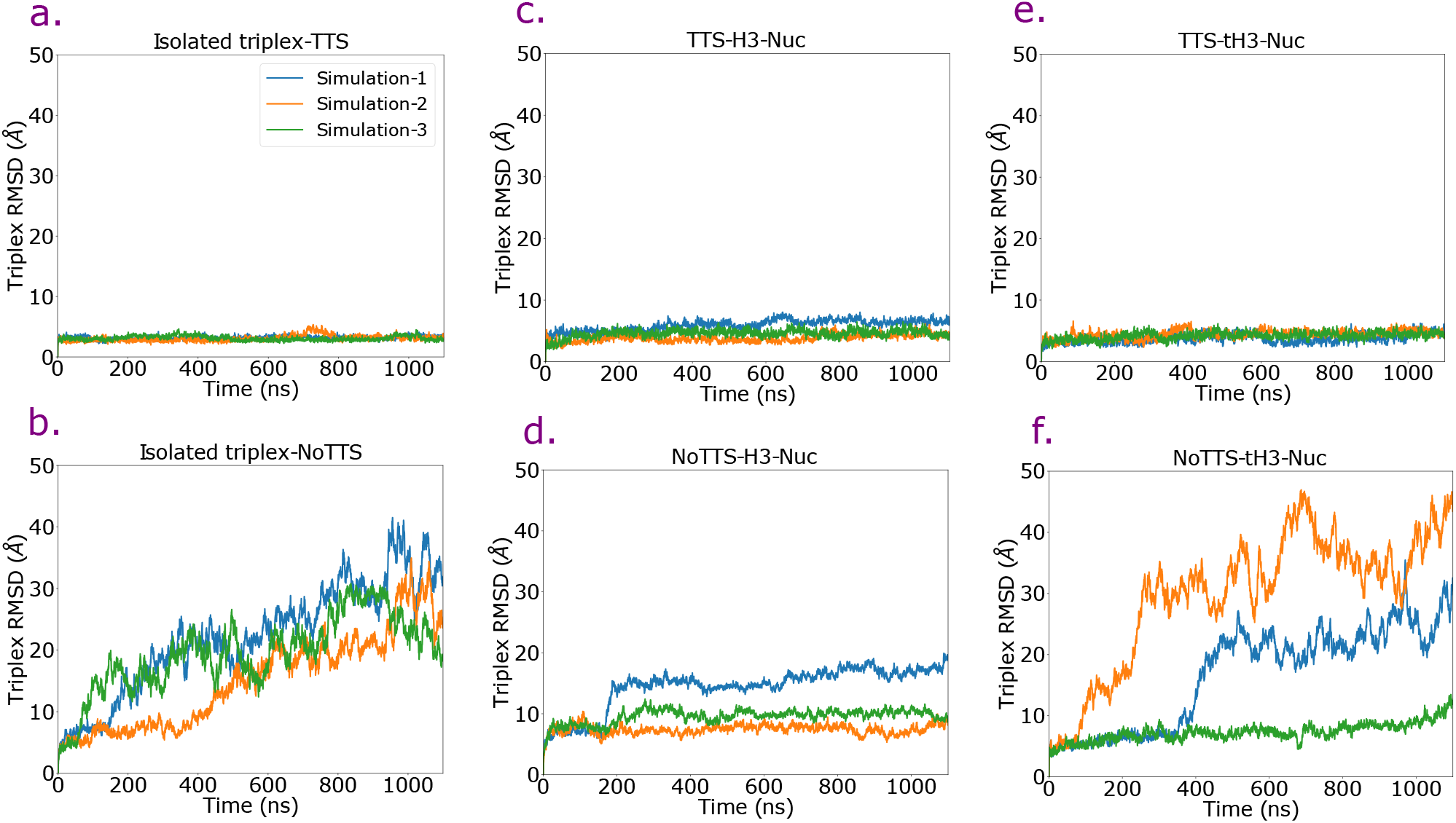
RMSD of RDD triplexes in isolated forms and in assembly with nucleosomes. The triplex dynamics are influenced by the DNA sequence at the triplex site. Models with the triplex targeting site sequence had the lowest RMSD values, while in samples with a NoTTS sequences the local and global stability of the RDD triplex was volatile and transient regardless of whether it was isolated or nucleosome bound.

These large RMSD values corresponded to the RNA detaching and unzipping from the DNA helix, as observed in snapshots of the final frames of the simulation trajectories (Fig 3d). A similar trend is noted by monitoring the RNA/DNA hydrogen-bond occupancies and the RNA/DNA distances (Figures 4 and S2). In all systems in which the DNA has a TTS sequence the number of RNA/DNA hydrogen bonds was well maintained beyond the terminal two residues, whereas the number of hydrogen bonds dropped dramatically in non-TTS systems. For non-TTS triplexes in complex with full-length nucleosomes there was additional hydrogen bond occupancy for the central basepairs 7-11 and 17-21, albeit not to the level as observed in TTS containing sequences. This partial localized stability was due to interactions with the H3 tail, as these additional hydrogen bonds were lost by removal of the H3 tail. The differences between the TTS and no-TTS systems is highly statistically significant, with p-values well below .001 (Table S2).

**Figure 3:**
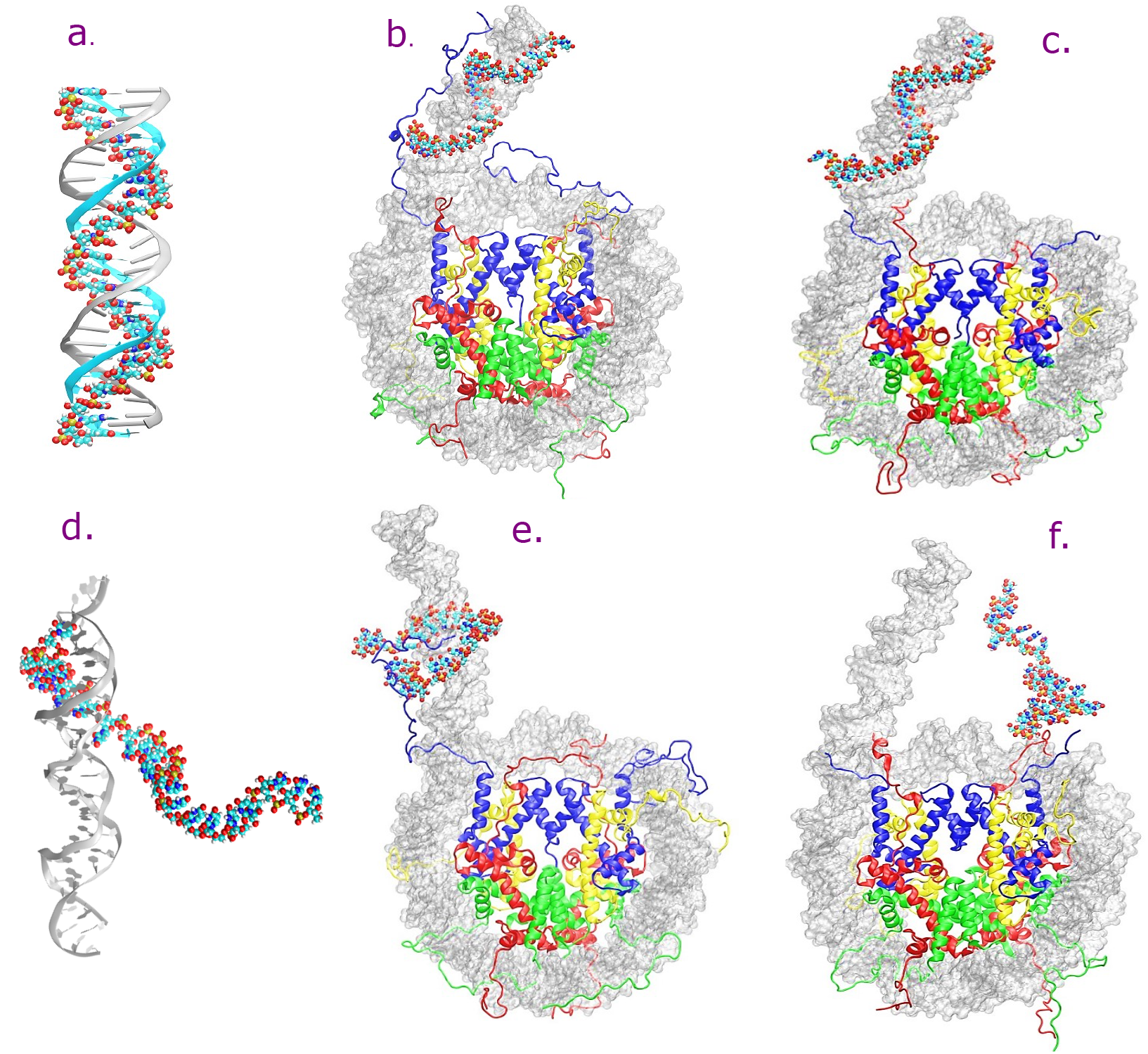
Final, representative snapshots from simulation trajectories. (a) Isolated RDD sequence with a triplex targeting site (TTS, in cyan) DNA sequence in cyan maintained its structural integrity. The RDD helix structure was maintained in nucleosome systems with (b) and without (c) the histone tails. For systems without the TTS sequence the RNA strand detached from the isolated DNA double helix (d), as well as the DNA in the full-length nucleosome (e) and the nucleosome lacking H3 tails (f).

**Figure 4:**
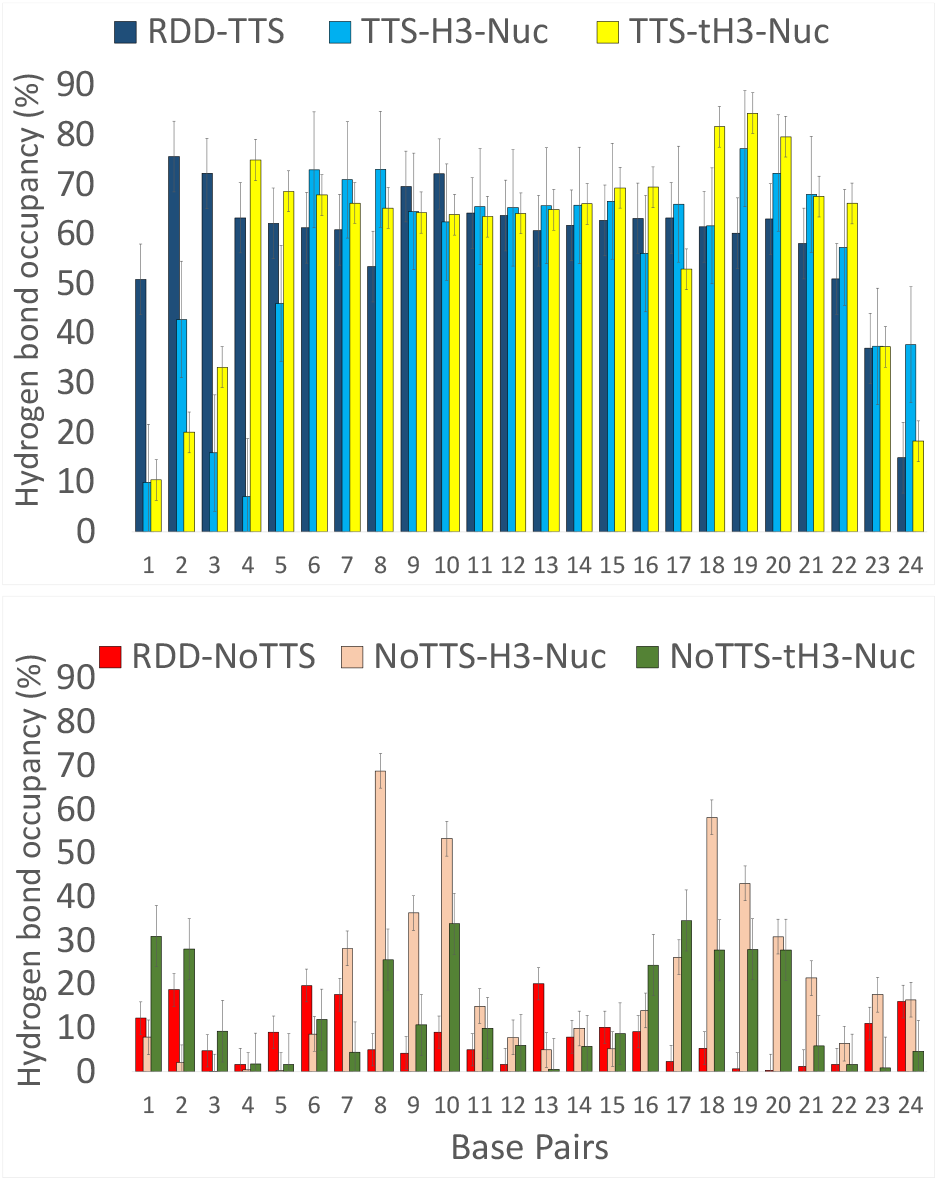
Occupancy of hydrogen bonds in RDD triplex between RNA and the adjacent DNA for systems containing a TTS DNA sequence (top) and lacking the TTS DNA sequence (bottom). The TFO sequence dictates the presence of two hydrogen bonds formed between U-A pairs in the RNA-DNA double helix. Table S2 shows the statistical differences between various models.

### 3.2 H3 tails form stronger interactions with RDD triplexes

In all simulations with the nucleosome, the protein core maintained its overall structure regardless of whether RNA was bound or not (Figure S3). The protein region with the most variability between structures were the H3 N-terminal tails. These regions are long, unstructured extensions which are key sites of multiple epigenetic markers. They are heavily charged, with four arginines and six lysines in the first 35 residues, and these extensions have been shown to influence nucleosome dynamics, DNA breathing, PTM accessibility, and effector protein binding (28, 59–67). Visual inspection of the simulation trajectories showed a dramatic difference in the H3 tail and entry DNA dynamics based on the presence or absence of RNA (Figures 3 and S7). This is in agreement with experiments in which the N-terminal tails of all histones, including H3, were truncated which showed a measurable reduction in the formation of RDD triplexes in the nucleosome (33). This reduced RDD stability in the absence of tails is likely a result of the lack of positively charged H3 residues in the tails at the DNA entry and exit sites of H3-truncated nucelosomes.

Given these key roles of the H3 tails in epigenetic regulation, we sought to identify how the H3 tails helped stabilize RNA binding, and to what degree RNA binding influenced the structure and dynamics of the H3 tails. In systems without RNA, the H3 tails at both the DNA entry and exit sites tended to form contacts with the DNA proximal to their location, which was primarily SHL 0.0-1.0 and -7.0 and the DNA arm for the entry DNA, and SHLs 6.0-7.0 for the exit DNA (Figures 1 and 5). In RNA and TTS containing systems, the entry DNA was more elongated along the DNA arm, with contacts primarily on the DNA arm and SHL -6.5 and -7.0. In addition, the exit DNA had increased contacts with the DNA near the dyad, with significantly more contacts formed to SHLs -1.5 to 0.5. The NoTTS case presented an intermediate of the two cases, with the entry H3 tail primarily having contacts with the DNA arm and SHL -7.0, whereas the exit H3 tail made some contacts with the dyad DNA, albeit fewer than in the case of the TTS containing sequence (Figure 5). This shifting of H3 tail contacts with the DNA by RNA is highlighted in a clustering analysis, which shows that in the TTS containing systems the dominant clusters have increased binding of the entry DNA along the DNA arm (Figure 6 and S4), whereas without RNA the H3 tail binding along the DNA arm is not the dominant structural cluster (Figure 6 and S5).

**Figure 5:**
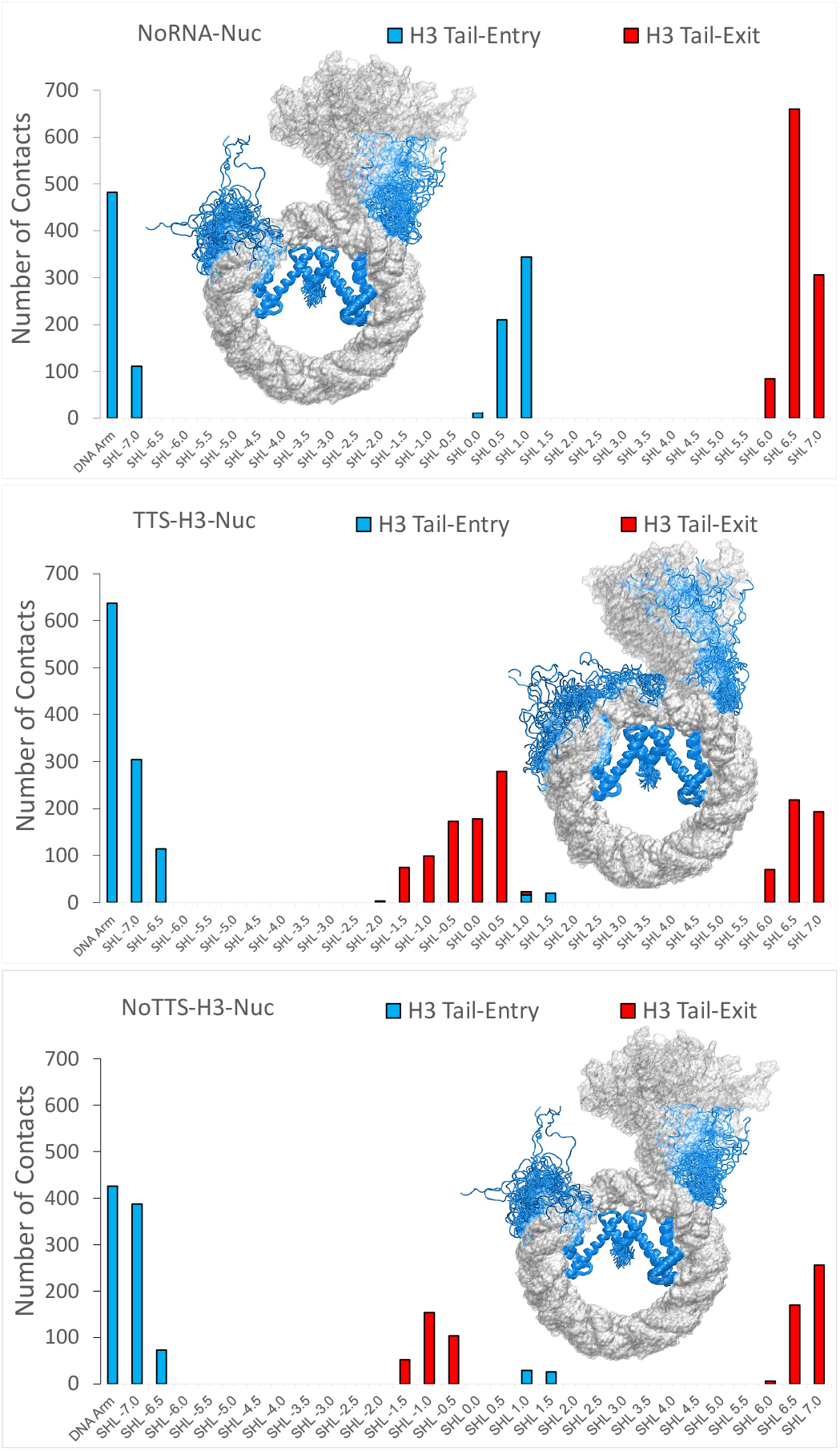
Superhelical locations of interactions between the H3 tails and nucleosomal DNA. The additional negative charge of RNA in the TTS and noTTS systems is attractive for both H3 tails at the entry and exit sites, resulting in a shift of interaction locations between the H3 tails and DNA.

**Figure 6:**
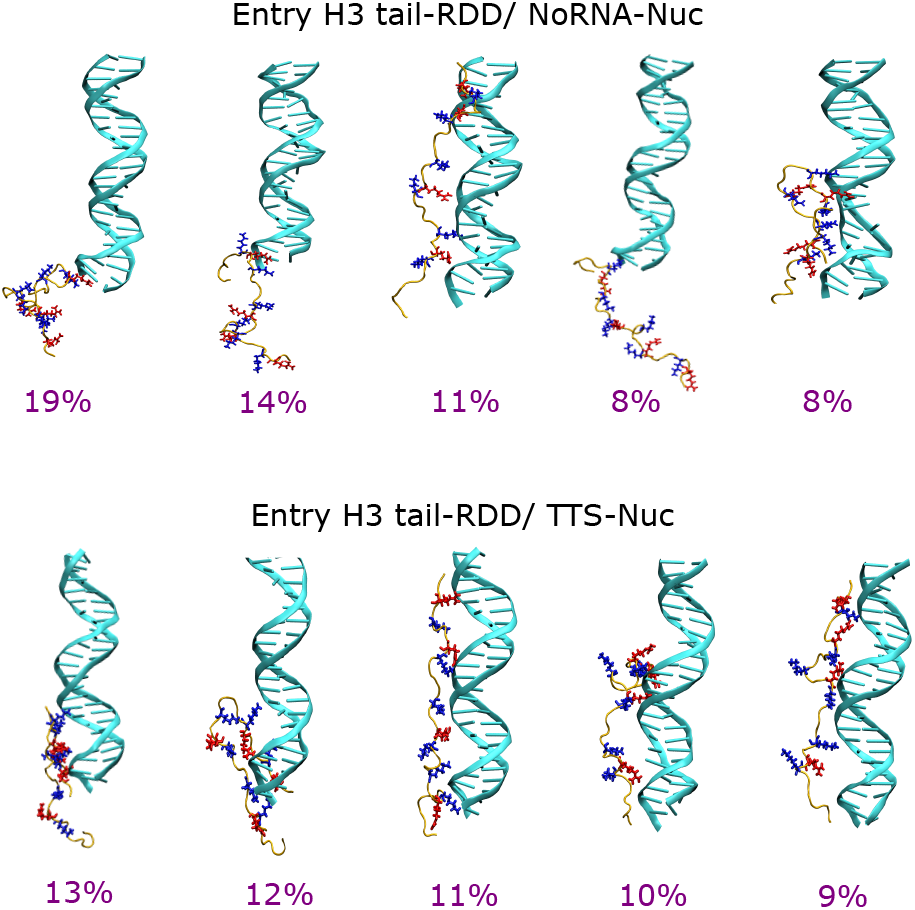
Top five dominant structural clusters of the H3 tails in the NoRNA and TTS nucleosomes. The introduction of a negatively charged RNA strand to nucleosomal DNA encourages the alignment of the H3 tail along the entry DNA arm in TTS nucleosomes.

An analysis of the hydrogen bond occupancy between the H3 tails and the nucleic acids shows a dramatic increase in hydrogen bonds upon RNA binding (Figure 7). In particular, for systems with RNA, hydrogen bond occupancies were significantly increased for Arg2, Arg8, Lys9, Arg17, and Arg26. This is likely due to the increased polyanionic properties of the RDD triplex, as its increased density of negative charges relative to the standard DNA double helix increase the attraction of and formation of hydrogen bonds to the H3 tails. These differences were found to be highly statistically significant (Table S3). In addition, while RNA presence helps to stabilize H3/nucleic acid interactions, the H3 tail presence also helps to stabilize RNA/DNA interactions. For systems which lacked the TTS sequence, absence of the H3 tail exacerbated disengagement of the RNA strand from the entry DNA arm (Figure 3). We also found that RNA binding affects the H3 tail flexibility and binding energy to the nucleic acids. The Root Mean Square Fluctuations (RMSFs) of the tails showed that for the entry H3 tail, systems with a TTS sequence had lower fluctuations in the first 10 residues, suggesting that tail binding was stabilized by interactions with the RDD triplex (Figures 8 and S6). The interaction energies between the entry and exit H3 histones and the nucleic acids were estimated through an MM/GBSA analysis, which primarily accounts for enthalpic terms in the interaction energies (Tables 1 and S4 -S6). Results show a significantly stronger binding of the entry H3 tails in systems with RNA, with an E_*total*_ for systems with RNA and with and without the TTS sequence of -80.23±1.65 kcal/mol and -77.74±1.5 kcal/mol compared to -52.15±0.99 kcal/mol in systems without RNA (Table 1). The energetics of the positively charged residues along the H3 tail interacting with the negatively charged nucleic acids varied significantly between systems (Tables S7 to S12). The most significant differences are associated with Arg2, Arg8, Arg17, Arg26 and Lys27, which had approximately doubled the affinity in TTS-nucleoeomes compared to NoRNA nucleoeomes. In particular, this interaction energy decreased by 3.98±1.01 kcal/mol and 2.38±1.12 kcal/mol respectively in TTS and NoTTS systems compared to the RNA free system in the case of Arg2, and for Arg8 there was a decrease of 6.27±0.83 kcal/mol and 5.07±0.9 kcal/mol for TTS and NoTTS nucleosomes compared to the NoRNA system (Tables S10 to S12). Notably, these residues are associated with multiple post-translational modifications which have been implicated in chromatin regulation (28, 63, 66, 68). A similar trend is observable for Lys4, Lys9, Arg17, Lys18, Lys23, Arg26, and Lys27 highlighting the influence of RNA on H3-DNA binding.

**Figure 7:**
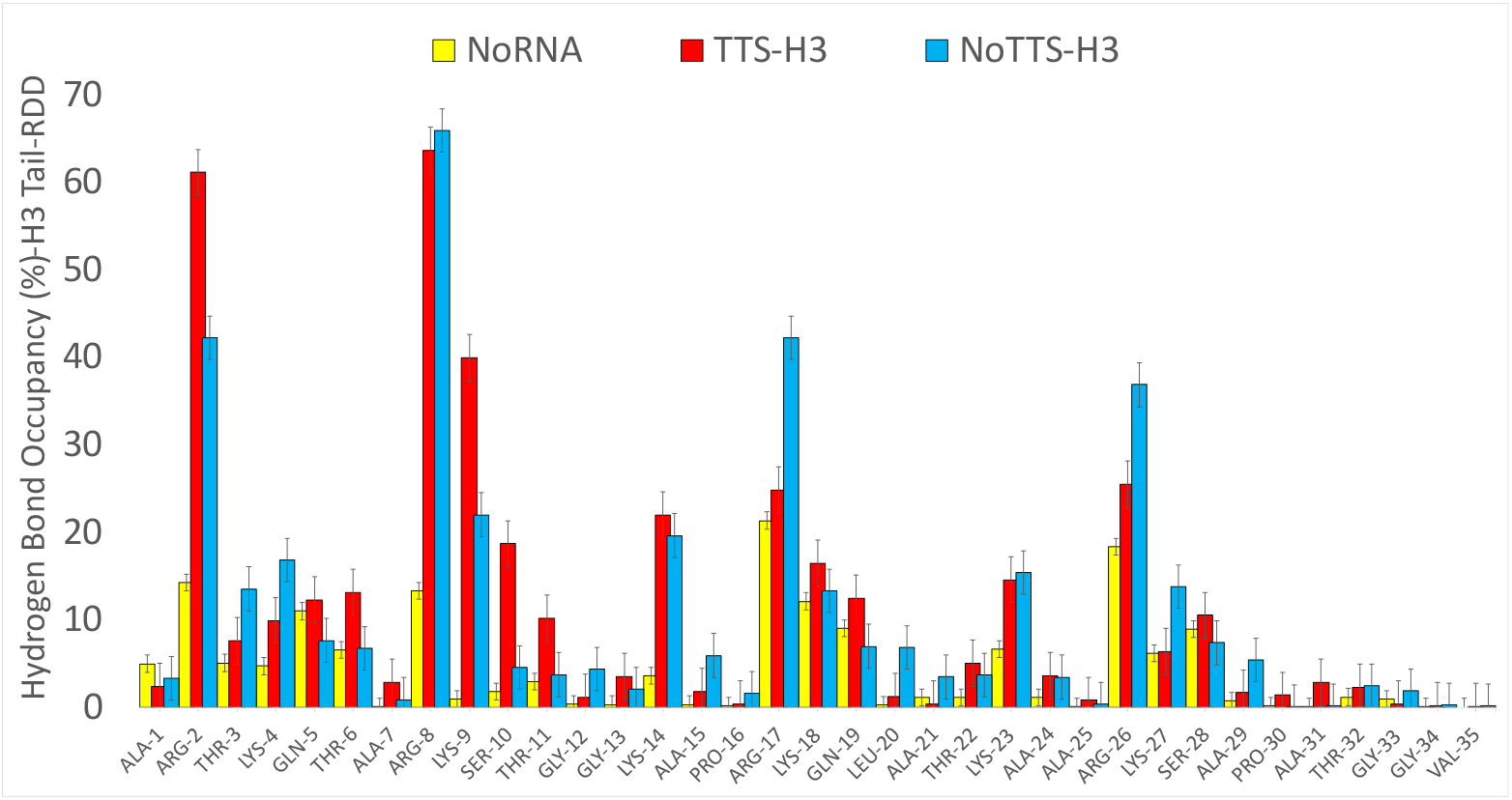
Average hydrogen bond occupancy formed between RDD and the adjacent H3 tails in each model. Arg2, Arg8, Lys9, Lys14, Arg17 and Arg26 of H3 tail are among the most favorable residues to form hydrogen bind with the nucleic acid triplex.

**Figure 8:**
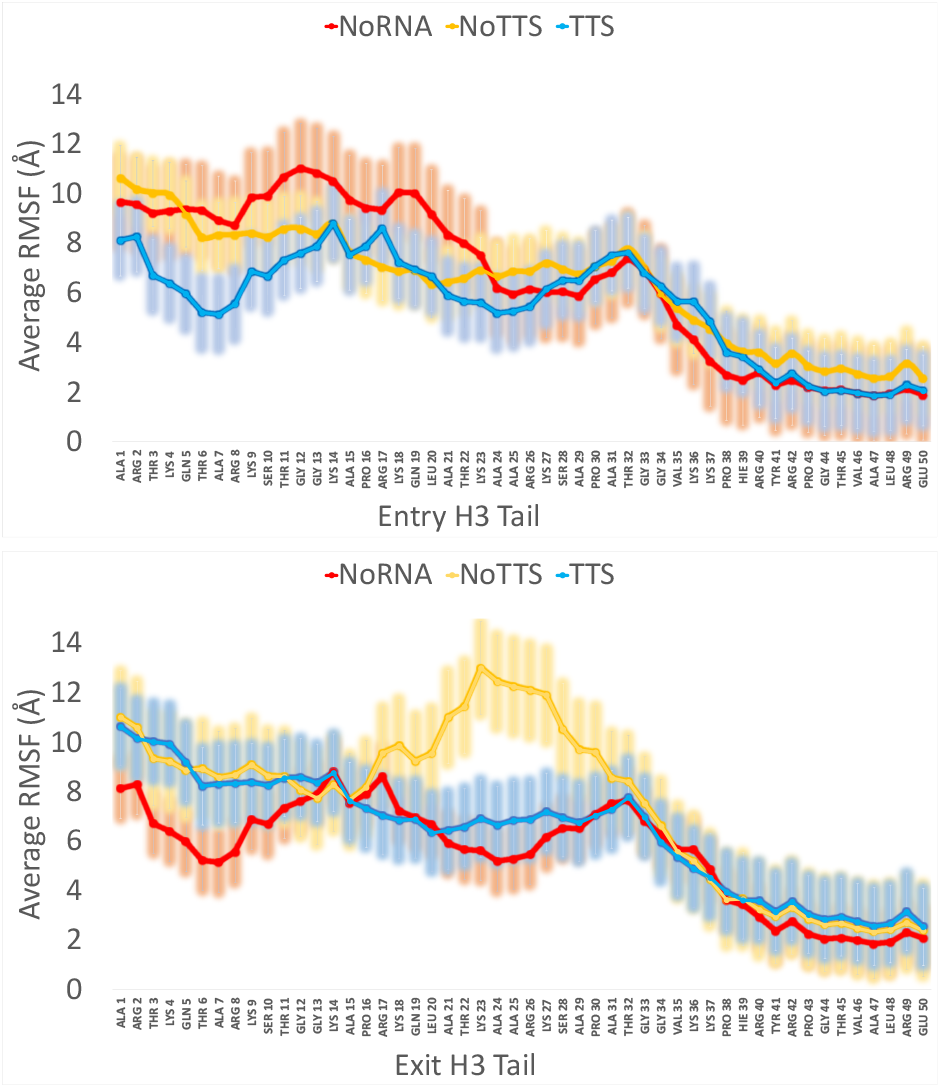
Average RMSF of the H3 tails at the entry (top) and exit (bottom) DNA sites show that RNA binding reduces tail flexibility in the H3 entry tail, particularly for the first 10 residues. Error bars represent standard error of mean.

**Table 1:**
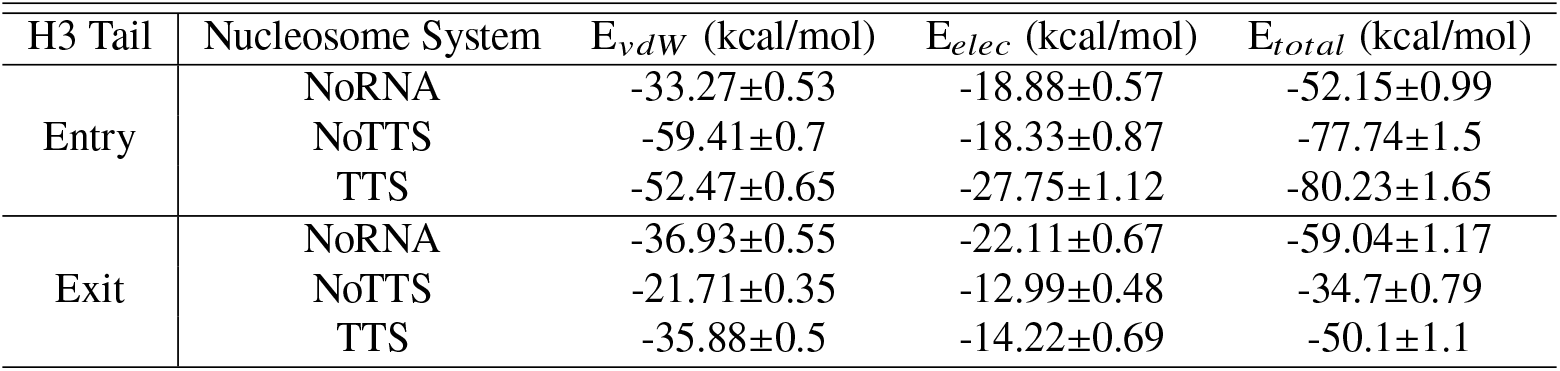
MM/GBSA interaction energies interaction (kcal/mol) between entry and exit H3 histone tails and nucleic acid (RNA-DNA) in nucleosome systems. Errors are reported as standard error of mean.

### 3.3 RNA influences the nucleosomal DNA structure

In addition to different H3 tail dynamics, RNA binding influenced the global and local structure of nucleosomal DNA. We did not observe any large-scale DNA unwrapping during our simulations, however introduction of RNA to the nucleosome did increases DNA breathing. More frequent DNA breathing is reflected in elevated entry-exit DNA end to end distances in the RNA included systems (Figures 9a and S8). The DNA entry-exit distances reached up to 120 Å in RNA-included nucleosomes, compared to only 90-100 Å in nucleosomes without RNA. We also observed reduction of the entry-exit DNA distance in nucleosomes lacking RNA to values as low as to 60 Å which signals the propensity of these systems to acquire a compact nucleosomal structure. In addition, the propensity for DNA opening in RNA-containing systems appears to depend on the presence of the H3 tails. For systems lacking the H3 tails there was an apparent decrease in the DNA end-to-end distances for the NoTTS system (Figure 9b), however this was primarily due to the dynamics in a single simulation and should therefore be interpreted with caution (Figure S8).

**Figure 9:**
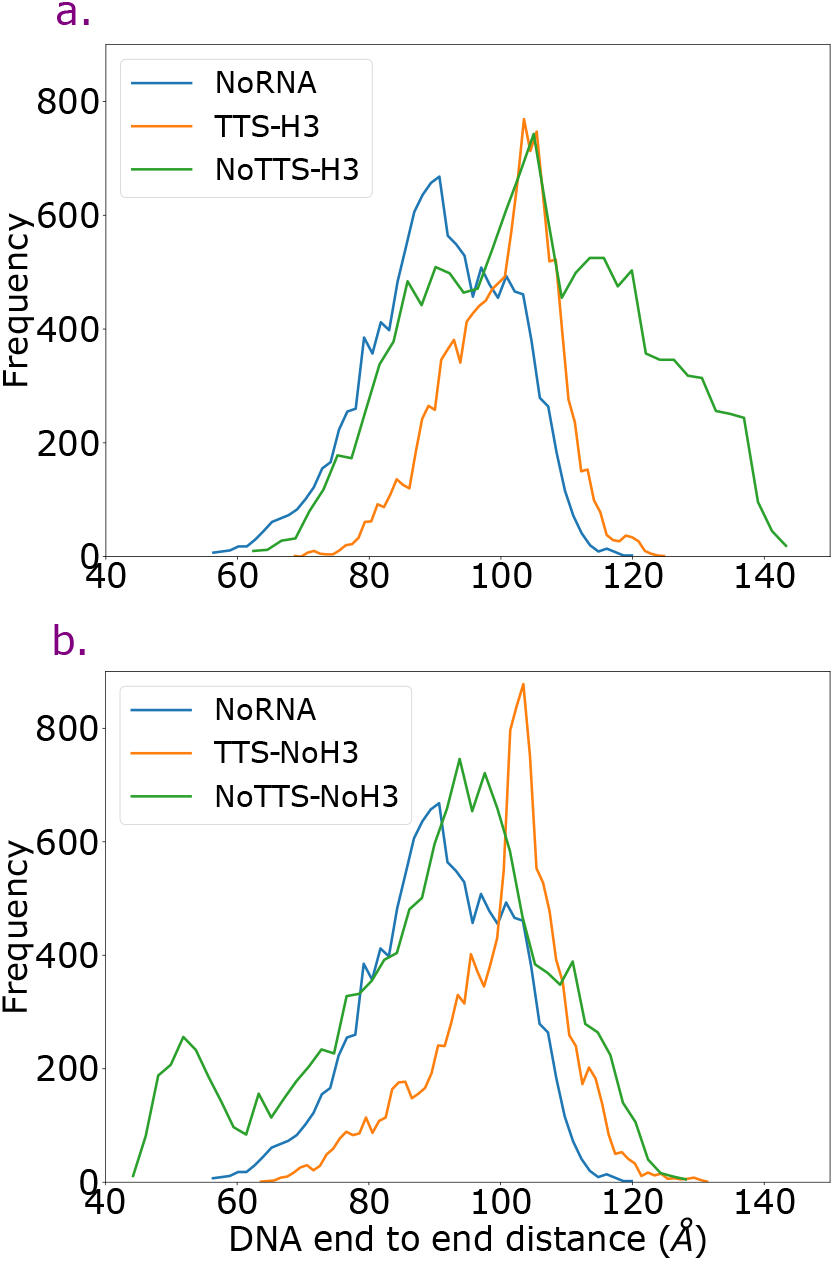
Histograms of DNA end to end distances for systems with H3 tails (top) and without H3 tails (bottom). For reference, the H3 tail containing noRNA system is shown in the bottom. The inclusion of RNA increased the distance distributions for systems with H3 tails, suggesting the propensity for increased DNA breathing.

Previous studies have suggested that the proper interaction of single-stranded RNA with a DNA double helix can result in the formation of a DNA triple helix with an intermediate structure between A and B-DNA forms (69). To examine how the DNA structures changes upon RNA binding in the context of the nucleosome, inter-base-pair nucleic acid parameters were computed for the DNA in the DNA arm from each simulation set. The averages of these values are summarized in Table S13 along with values for ideal A-and B-DNA structures (69). From these six metrics, slide and shift showed the most deviation from ideal B-DNA towards ideal A-DNA (Figures 10, S9-S14). Models without the TTS sequence showed the highest bias to the A-DNA roll value, while all models show negative slide. The combination of negative slide and positive roll are hallmarks of A-DNA (70). In contrast, models with a TTS sequence had slide values close to the A-DNA state. Although the range of change for the base-pair shift is narrow (the shift for ideal B-DNA is 0.0 Å and for A-DNA is 0.01 Å) all models showed an increasing trend for this parameter. The basepair step inclination and X-displacement are the major geometrical parameters that track the global structural changes in the DNA double helix (71). These parameters are zero for an ideal B-DNA duplex, while for the case of ideal A-DNA they are 20° and -5 Å respectively (Table S14). Elevated values of the helix X-displacement and inclination were observed in all TTS and NoTTS systems, indicating DNA structural variations from linear B-DNA structures towards structures with global shapes resembling A-form (Figures 10 & S15-S16). We also observed a strong correlation between variations of local base-pair and and helix parameters in all TTS and NoTTS systems (Figures S17 & S18). These correlations indicate that angular and linear variations in base-pair coordinates have a cumulative influence on the DNA helix structure which is invariant under sequence type or the presence or absence of RNA and H3 tails.

**Figure 10:**
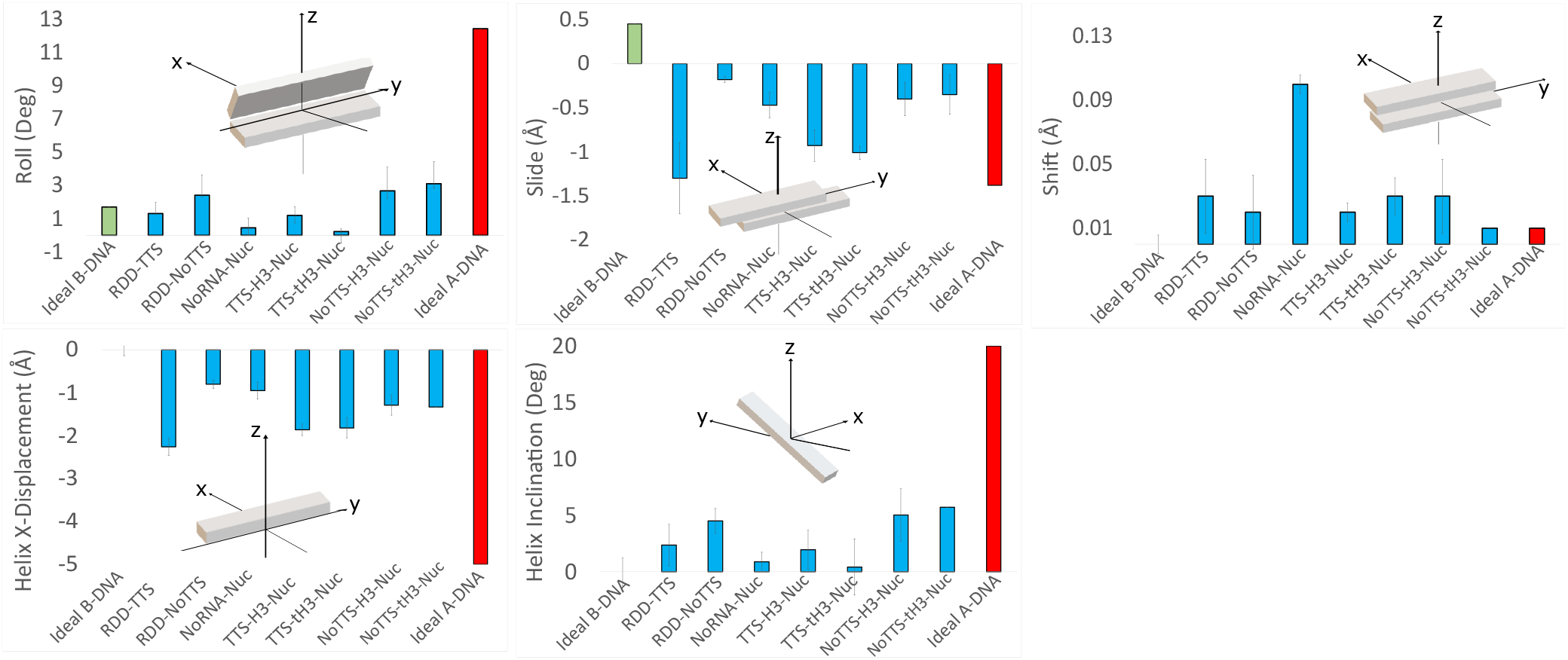
Deviations from ideal A and B-DNA values for inter base-pair parameters of roll, slide and shift and base-pair step helix parameters of x-displacement and inclination. Base-pair step helix inclination, X-displacement and inter base-pair shift values are zero for ideal B-DNA.

## 4 DISCUSSION

The binding of small ncRNAs to nucleosomal DNA triggers a sequence of events that modifies the structure and energetics of both the DNA and the critical N-terminal H3 tails. Simulations show that the stability of an RNA.DNA-DNA triple helix is highly-sequence dependent, as even in the context of a nucleosome with full-length H3 tails RNA separation was observed with a non-complimentary DNA sequence on the hundreds of nanosecond timescale. Stable RNA binding increases the attraction of the H3 tails with the nucleic acids, as shown by both the increased hydrogen binding and decreased MM/GBSA-derived energies computed here. This effect appears to be largely due to the increased acidic nature of the triple helix, which creates a strong electrostatic attraction for the multiple arginines and lysines in the tails. RDD triplex formation also has the effect of altering the DNA structure to a shape that is intermediate of ideal B-and A-forms.

The effects of RNA binding on the H3 tails likely has significant effects on gene regulation. Over a third of the residues in these tails have been identified as sites for post-translational modifications that are recognized by various reader proteins (68). Simulations and NMR experiments have shown that the energetics and dynamics of these tails on nucleosomal DNA can be tuned by reducing the charge states of specific residues, for example through acetylation or phosphorylation, and that by doing so other epigentic markers may become more accessible to reader proteins (28). This provides one example of cross-talk in which one epigenetic factor influences another at a distant location (72). Our results suggest RNA binding may have the opposite effect. By forming a stronger H3/nucleic acid interface, PTM locations used for various signaling pathways are likely more occluded than in systems without RNA, making the addition and reading of some PTMs more difficult. Given that RNA binding and TTSs are enriched at regulatory elements that correlate with active chromatin locations in vivo (33), the interplay between RNA and other epigenetic marks, particularly those that utilize the H3 tails, is likely to have implications for highly expressed genes.

Our results also show how RNA binding influences global and local DNA structures. Although these simulations were performed on only single nucleosome systems, we can infer that these changes may also influence higher-order chromatin structures. Indeed, cryo-EM and crystal structures have shown that heterogeneous poly-nucleosomal arrays can form regular structures in solution (73–75), while simulations have shown that these structures are dynamic in nature and can be modulated by factors such as linker histone binding and linker DNA length (76–79). The introduction of heterogeneity to these structures, such as through subtle changes to the nucleosomal DNA structure or by changing the linker DNA length, can disrupt these higher-order structures (80, 81). As shown here, ncRNA binding influences local and global DNA structures. These changes likely create another source of conformational heterogeneity in chromatin fibers, thus disrupting the repeating structures observed in some experiments. Additional experiments and simulations with ncRNA binding will, therefore, likely show a loss of ordering from regularly repeating poly-nucleosomal arrays upon the introduction of ncRNAs.

## Supporting information

Supporing Information

## ACKNOWLEDGMENTS

The authors thank members of the Wereszczynski Group for valuable discussions concerning this work. This work used the Extreme Science and Engineering Discovery Environment (XSEDE) (82), This research was supported by the National Institute of General Medical Sciences of the National Institutes of Health Grant R35GM119647. The content is solely the responsibility of the authors and does not necessarily represent the official views of the National Institutes of Health.

