## Supplementary material for "The Effects of RNA.DNA-DNA Triple Helices on Nucleosome Structures and Dynamics": Supporing Information

### Supporting Information for The Effects of RNA•DNA-DNA Triple Helices on Nucleosome Structures and Dynamics

Table S1: A summary of structures and molecular dynamics simulations performed in this study.

| System | Number of Atoms | Number of Replica | Time | Total |
|---|---|---|---|---|
| Isolated Triplex TTS | 2253 | 3 | 1.1 µs | 3.3 µs |
| Isolated Triplex NoTTS | 2240 | 3 | 1.1 µs | 3.3 µs |
| NoRNA-Nuc | 26244 | 3 | 1.1 µs | 3.3 µs |
| TTS-Nuc$^{tH3}$ | 25897 | 3 | 1.1 µs | 3.3 µs |
| TTS-Nuc | 26965 | 3 | 1.1 µs | 3.3 µs |
| NoTTS-Nuc$^{tH3}$ | 25884 | 3 | 1.1 µs | 3.3 µs |
| NoTTS-Nuc | 26952 | 3 | 1.1 µs | 3.3 µs |

Table S2: There is a significant statistical difference between TTS and NoTTS models for the RNA/DNA hydrogen bond occupancies.

| p-Value | RDD-NoTTS | TTS-Nuc | TTS-Nuc$^{tH3}$ | NoTTS-Nuc | NoTTS-Nuc$^{tH3}$ |
|---|---|---|---|---|---|
| RDD-TTS | **1.49E-22** | 0.4 | 0.95 | **8.50E-11** | **9.90E-17** |
| RDD-NoTTS | | **2.57E-14** | **1.46E-15** | **6.00E-03** | **3.00E-02** |
| TTS-Nuc | | | 0.5 | **1.76E-07** | **5.52E-11** |
| TTS-Nuc$^{tH3}$ | | | | **1.61E-08** | **3.44E-12** |
| NoTTS-Nuc | | | | | 2.20E-01 |

Table S3: Statistical difference for the number of hydrogen bonds between the H3 tails and RDD triplexes.

| p-Value | TTS-Nuc | NoTTS-Nuc |
|---|---|---|
| NoRNA-Nuc | **7.50E-03** | **7.10E-03** |
| TTS-Nuc | | 0.99 |

Table S4: MM/GBSA interaction energies (kcal/mol) between H3 histones and DNA.

| Nucleosome System | $E_{vdW}$ | $E_{elec}$ | $E_{total}$ |
|---|---|---|---|
| NoRNA | -254.93±3.3 | -176.98± 4.12 | -431.91±3.52 |
| NoTTS | -264.8±1.91 | -154.28±2.24 | -419.09±3.38 |
| TTS | -270.54±1.9 | -165.87±2.36 | -436.42±3.52 |

Table S5: MM/GBSA interaction energies (kcal/mol) between the exit H3 histone and DNA.

| Nucleosome System | $E_{vdW}$ | $E_{elec}$ | $E_{total}$ |
|---|---|---|---|
| NoRNA | -125.12±0.93 | -76.58±1.02 | -201.71±1.59 |
| NoTTS | -109.19±0.92 | -66.07±1.05 | -175.26±1.54 |
| TTS | -120.86±0.92 | -66.16±1.02 | -187.03±1.58 |

Table S6: MM/GBSA interaction energies (kcal/mol) between the entry H3 histone and DNA.

| Nucleosome System | $E_{vdW}$ | $E_{elec}$ | $E_{total}$ |
|---|---|---|---|
| NoRNA | -129.81±0.97 | -100.4±1.36 | -230.21±1.93 |
| NoTTS | -155.61±0.99 | -88.21±1.24 | -243.83±1.84 |
| TTS | -149.68±0.98 | -99.71±1.34 | -249.39±1.94 |

Table S7: Residue-wise decomposition of MM/GBSA interaction energies between the H3 tails and nucleic acids in the no RNA system (kcal/mol).

| Residue | $E_{vdW}$ | $E_{elec}$ | $E_{total}$ |
|---|---|---|---|
| ARG 2 | -4.23±2 | -1.53±0.5 | -5.76±1.25 |
| ARG 8 | -1.92±0.92 | -3.49±0.15 | -5.41±0.53 |
| ARG 17 | -2.20±0.98 | -3.13±0.32 | -5.34±0.65 |
| ARG 26 | -3.39±1.49 | -1.03±0.13 | -4.42±0.81 |
| LYS 4 | -1.66±0.76 | -2.25±0.47 | -3.90±0.61 |
| GLN 5 | -1.98±0.82 | -1.43±0.57 | -3.41±0.7 |
| THR 6 | -0.77±0.47 | -1.91±0.68 | -2.68±0.57 |
| LYS 18 | -1.17±0.69 | -1.40±0.55 | -2.57±0.62 |
| THR 32 | -0.92±0.57 | -1.54±0.68 | -2.45±0.63 |
| LYS 23 | -0.98±0.65 | -1.04±0.23 | -2.03±0.44 |
| LYS 27 | -2.03±0.91 | 0.20±0.3 | -1.83±0.6 |
| SER 28 | -0.92±0.63 | -0.75±0.32 | -1.67±0.48 |
| ALA 1 | -1.43±0.54 | -0.09±0.16 | -1.52±0.35 |
| LYS 9 | -0.76±0.55 | -0.70±0.21 | -1.46±0.38 |
| ALA 31 | -0.88±0.53 | -0.26±0.36 | -1.14±0.45 |

Table S8: Residue-wise decomposition of MM/GBSA interaction energies between the H3 tails and nucleic acids in the noTTS system (kcal/mol).

| Residue | $E_{vdW}$ | $E_{elec}$ | $E_{total}$ |
|---|---|---|---|
| ARG 8 | -4.97±1.42 | -5.51±1.1 | -10.48±1.26 |
| ARG 26 | -4.64±1.81 | -4.24±0.21 | -8.88±1.01 |
| ARG 2 | -3.78±1.2 | -4.37±0.79 | -8.14±1 |
| ARG 17 | -4.59±1.06 | -2.74±0.78 | -7.33±0.92 |
| LYS 27 | -1.86±0.89 | -1.98±0.95 | -3.85±0.92 |
| LYS 9 | -2.94±0.86 | -0.87±0.36 | -3.81±0.61 |
| LYS 14 | -2.99±1.27 | -0.19±0.26 | -3.18±0.77 |
| LYS 4 | -2.14±1.18 | -1.01±0.55 | -3.15±0.87 |
| LYS 23 | -1.64±0.77 | -1.41±0.08 | -3.05±0.42 |
| LYS 18 | -1.61±0.67 | -1.10±0.7 | -2.71±0.69 |
| LEU 20 | -2.43±1.01 | 0.33±0.05 | -2.10±0.53 |
| SER 28 | -0.87±0.48 | -0.91±0.65 | -1.78±0.57 |
| THR 3 | -1.18±0.63 | -0.56±0.52 | -1.74±0.58 |
| ALA 21 | -1.44±0.71 | 0.12±0.04 | -1.32±0.38 |
| GLN 5 | -1.95±0.81 | 0.66±0.2 | -1.29±0.51 |

Table S9: Residue-wise decomposition of MM/GBSA interaction energies between the H3 tails and nucleic acids in the TTS system (kcal/mol).

| Residue | $E_{vdW}$ | $E_{elec}$ | $E_{total}$ |
|---|---|---|---|
| ARG 8 | -3.42±0.84 | -8.26±1.43 | -11.67±1.13 |
| ARG 2 | -3.54±1.27 | -6.2±0.28 | -9.74±0.78 |
| LYS 9 | -3.4±0.85 | -4.22±0.65 | -7.62±0.75 |
| ARG 26 | -4.23±1.33 | -2.56±0.11 | -6.79±0.72 |
| ARG 17 | -4.24±2.02 | -2.33±0.84 | -6.57±1.43 |
| LYS 14 | -1.88±0.95 | -2.59±0.84 | -4.46±0.9 |
| LYS 23 | -2.38±0.98 | -0.96±0.01 | -3.8±0.5 |
| LYS 18 | -2.19±0.8 | -1.47±0.38 | -3.68±0.59 |
| LYS 4 | -1.61±0.79 | -1.17±0.26 | -2.78±0.53 |
| LYS 27 | -1.82±1.2 | -0.15±0.06 | -1.97±0.63 |
| GLN 5 | -2.55±1.07 | 0.83±0.3 | -1.72±0.68 |
| SER 10 | -0.74±0.44 | -0.92±0.64 | -1.67±0.54 |
| LEU 20 | -1.73±1.34 | 0.2±0.05 | -1.53±0.69 |
| THR 6 | -1.03±0.48 | -0.44±0.3 | -1.48±0.39 |
| PRO 30 | -2.01±1.19 | 0.55±0.05 | -1.46±0.62 |

Table S10: Significant differences in the interaction energies for the H3 tail residues between the NoRNA and TTS systems (kcal/mol). Positive values represent more favorable interactions in the TTS system.

| Residue | $\Delta E_{vdW}$ | $\Delta E_{elec}$ | $\Delta E_{total}$ |
|---|---|---|---|
| ARG 2 | -0.69±1.63 | 4.67±0.39 | 3.98±1.01 |
| LYS 4 | -0.05±0.78 | -1.07±0.36 | -1.12±0.57 |
| ARG 8 | 1.50±0.88 | 4.77±0.79 | 6.27±0.83 |
| LYS 9 | 2.64±0.7 | 3.52±0.43 | 6.17±0.57 |
| ARG 17 | 2.04±1.5 | -0.80±0.58 | 1.24±1.04 |
| LYS 18 | 1.02±0.75 | 0.07±0.47 | 1.09±0.61 |
| LYS 23 | 1.85±0.82 | -0.08±0.12 | 1.77±0.47 |
| ARG 26 | 0.84±1.41 | 1.53±0.12 | 2.38±0.76 |
| LYS 27 | -0.21±1.06 | 0.34±0.18 | 0.13±0.62 |

Table S11: Significant differences in the interaction energies for the H3 tail residues between the NoRNA and noTTS systems (kcal/mol). Positive values represent more favorable interactions in the noTTS system.

| Residue | $\Delta E_{vdW}$ | $\Delta E_{elec}$ | $\Delta E_{total}$ |
|---|---|---|---|
| ARG 2 | -0.46±1.6 | 2.84±0.65 | 2.38±1.12 |
| LYS 4 | 0.48±0.97 | -1.24±0.51 | -0.75±0.74 |
| ARG 8 | 3.05±1.17 | 2.02±0.62 | 5.07±0.9 |
| LYS 9 | 2.18±0.71 | 0.17±0.29 | 2.35±0.5 |
| ARG 17 | 2.39±1.02 | -0.39±0.55 | 2±0.79 |
| LYS 18 | 0.43±0.68 | -0.3±0.62 | 0.13±0.65 |
| LYS 23 | 0.66±0.71 | 0.36±0.15 | 1.02±0.43 |
| ARG 26 | 1.25±1.65 | 3.21±0.17 | 4.46±0.91 |
| LYS 27 | -0.17±0.9 | 2.18±0.62 | 2.01±0.76 |

Table S12: Differences in the energy of interaction for common H3 residues between TTS and NoTTS nucleosomes. Positive values are favorable in TTS, negative are favorable in noTTS. Energies in kcal/mol.

| Residue | $\Delta E_{vdW}$ | $\Delta E_{elec}$ | $\Delta E_{total}$ |
|---|---|---|---|
| ARG 2 | -0.23±1.24 | 1.83±0.54 | 1.6±0.89 |
| LYS 4 | -0.53±0.99 | 0.17±0.41 | -0.37±0.7 |
| ARG 8 | -1.55±1.13 | 2.75±1.26 | 1.2±1.2 |
| LYS 9 | 0.46±0.86 | 3.35±0.5 | 3.81±0.68 |
| ARG 17 | -0.35±1.54 | -0.41±0.81 | -0.76±1.17 |
| LYS 18 | 0.59±0.74 | 0.37±0.54 | 0.96±0.64 |
| LYS 23 | 1.19±0.87 | -0.44±0.04 | 0.75±0.46 |
| ARG 26 | -0.41±1.57 | -1.68±0.16 | -2.08±0.86 |
| LYS 27 | -0.04±1.04 | -1.84±0.5 | -1.88±0.77 |

Table S13: Deviation from ideal B and A-DNA inter base-pair parameters of tilt, twist, roll, slide, shift and rise.

| System | Tilt (Deg) | Twist (Deg) | Roll (Deg) | Slide (Å) | Shift (Å) | Rise (Å) |
|---|---|---|---|---|---|---|
| Ideal B-DNA | ≈ -0.01 | ≈ 35.96 | ≈ 1.71 | ≈ 0.45 | ≈ 0.0 | ≈ 3.36 |
| Ideal A-DNA | ≈ -0.08 | ≈ 30.31 | ≈ 12.43 | ≈ -1.38 | ≈ 0.01 | ≈ 3.3 |
| RDD-TTS | 0.06 ± 0.12 | 32.18 ± 5.26 | 1.32 ± 0.66 | -1.3 ± 0.4 | 0.03 ± 0.01 | 3.21 ± 0.03 |
| RDD-NoTTS | 0.17 ± 0.12 | 32.42 ± 9.75 | 2.42 ± 1.22 | -0.18 ± 0.03 | 0.02 ± 0.02 | 3.16 ± 0.03 |
| NoRNA-Nuc | 0.55 ± 0.09 | 34.67 ± 9.65 | 0.46 ± 0.58 | -0.47 ± 0.14 | 0.1 ± 0.02 | 3.3 ± 0.07 |
| TTS-Nuc | 0.58 ± 0.31 | 30.27 ± 6.41 | 1.21 ± 0.5 | -0.93 ± 0.18 | 0.02 ± 0.01 | 3 ± 0.23 |
| TTS-Nuc$^{tH3}$ | -0.1 ± 0.09 | 33.17 ± 9.66 | 0.23 ± 0.16 | -1.01 ± 0.08 | 0.03 ± 0.01 | 3.24 ± 0.4 |
| NoTTS-Nuc | 0.04 ± 0.09 | 31.58 ± 9.32 | 2.67 ± 1.44 | -0.4 ± 0.19 | 0.03 ± 0.02 | 3.16 ± 0.27 |
| NoTTS-Nuc$^{tH3}$ | 0.19 ± 0.12 | 31.93 ± 7.84 | 3.1 ± 1.33 | -0.35 ± 0.23 | 0.01 ± 0.04 | 3.23 ± 0.16 |

Table S14: Deviation from ideal B and A-DNA base-pair step helix parameters.

| System | Inclination (Deg) | X-displacement (Å) |
|---|---|---|
| Ideal B-DNA | ≈ 0.0 | ≈ 0.0 |
| Ideal A-DNA | ≈ 20 | ≈ -5.0 |
| RDD-TTS | 2.37 ± 1.21 | -2.26 ± 0.13 |
| RDD-NoTTS | 4.52 ± 1.85 | -0.8 ± 0.2 |
| NoRNA-Nuc | 0.88 ± 1.1 | -0.95 ± 0.1 |
| TTS-Nuc | 1.96 ± 0.87 | -1.86 ± 0.2 |
| TTS-Nuc$^{tH3}$ | 0.43 ± 1.73 | -1.82 ± 0.13 |
| NoTTS-Nuc | 5.05 ± 2.48 | -1.29 ± 0.23 |
| NoTTS-Nuc$^{tH3}$ | 5.73 ± 2.31 | -1.33 ± 0.23 |

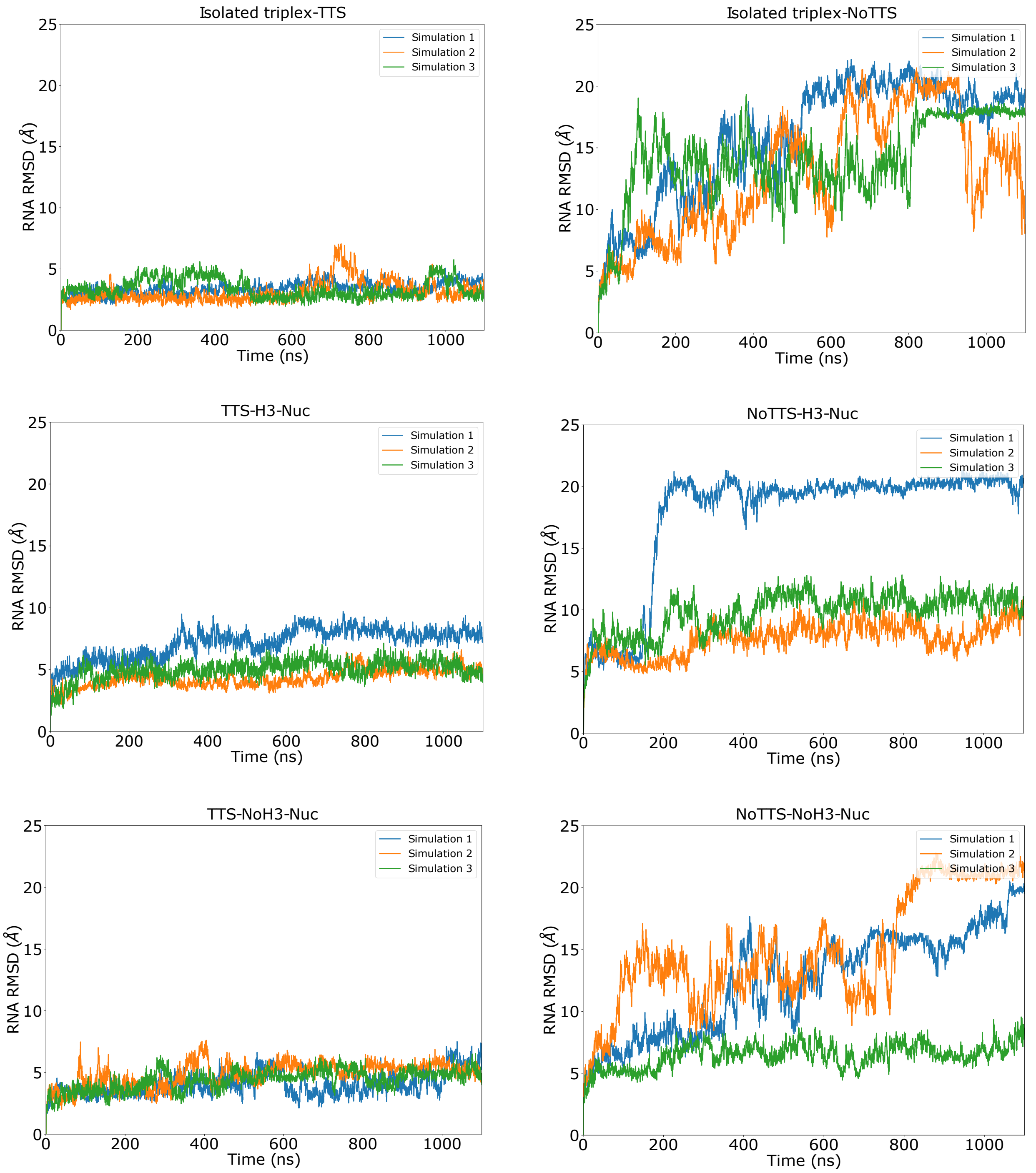


Figure S1: RMSD of RNA in isolated triplex and nucleosome systems. RNA is the most volatile and highly dynamic segment of the RDD triple helix. RMSD of RNA can reach up to 23 Å in samples that do not contain TTS, showing that DNA sequence and its compatibility with triplex-forming oligonucleotide plays a major role in robust binding of RNA and DNA at the entry/exit sites of nucleosomes.

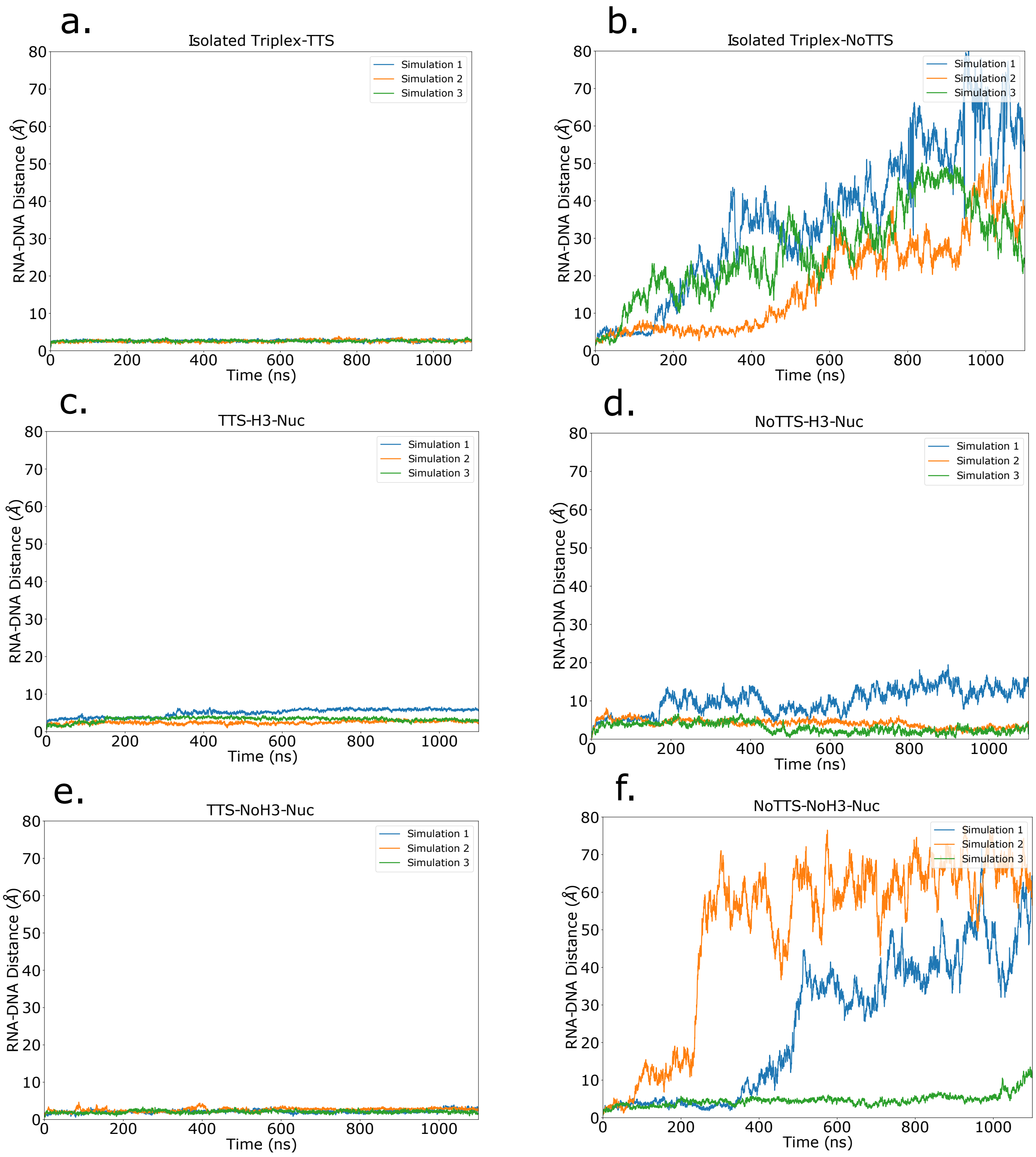


Figure S2: DNA and RNA distance in isolated and nucleosome systems. In the absence of the TTS sequence, RNA is released from its adjacent DNA double helix and the sequence dependent event is observed in isolated NoTTS triplex and NoTTS-nucleosomes$^{tH3}$s.

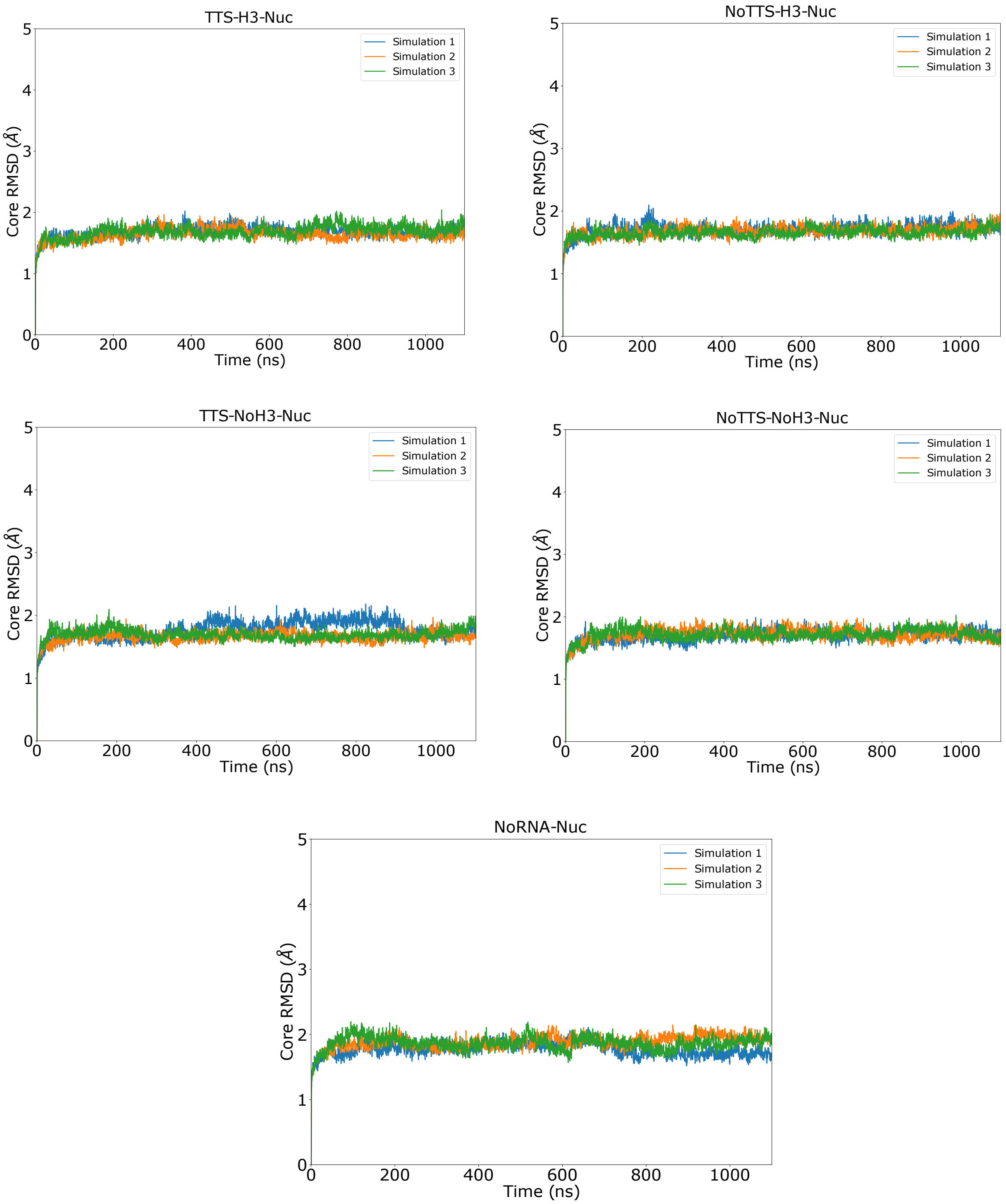


Figure S3: RMSD of the histone protein core in nucleosomal systems. Addition of RNA to the nucleosome does not effect the secondary structure of the major histone α-helices.

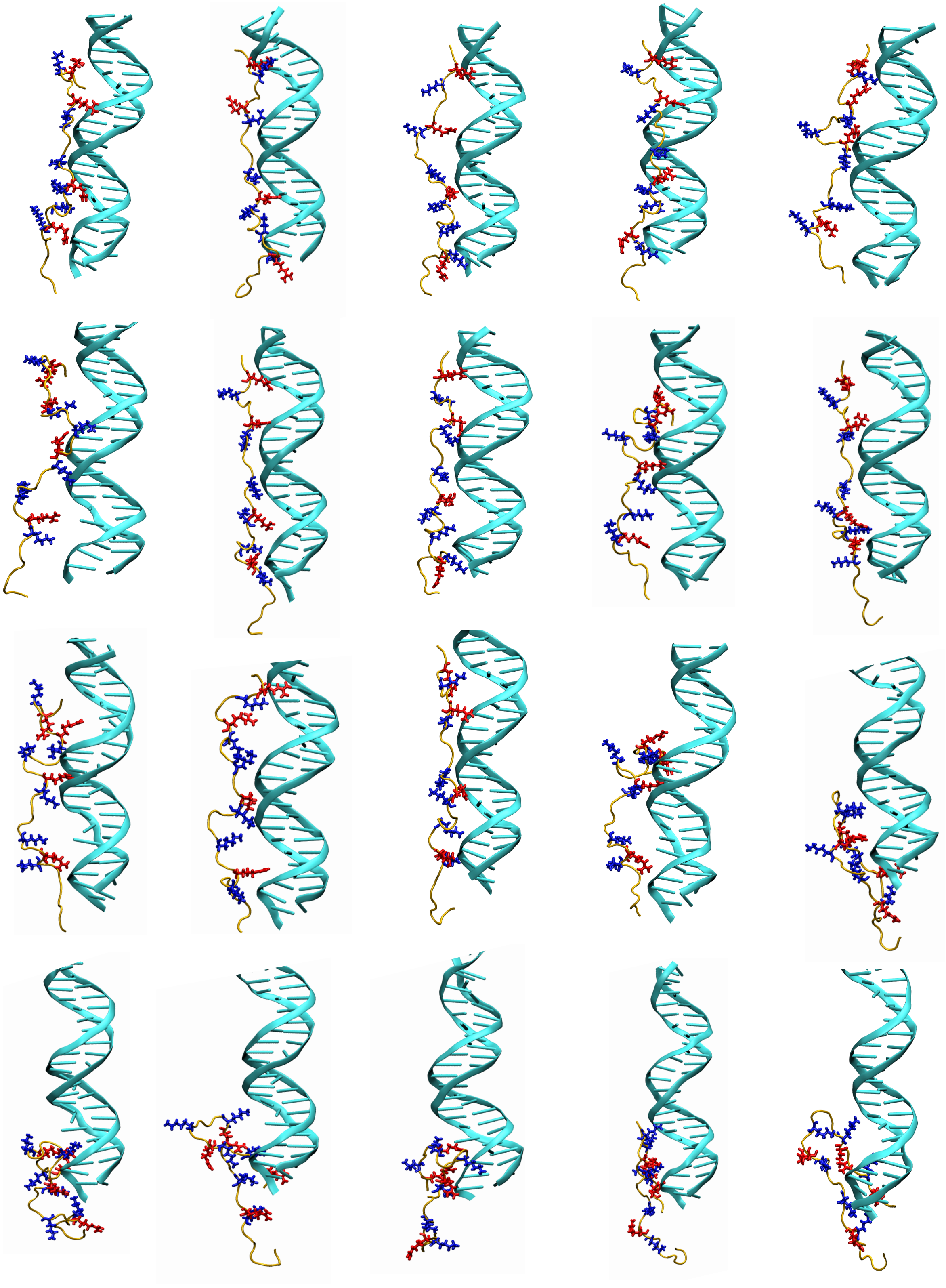

Figure S4: Introduction of RNA to entry DNA in TTS nucleosomes enhances the alignment of long arginine and lysine amino acids with DNA major grooves.

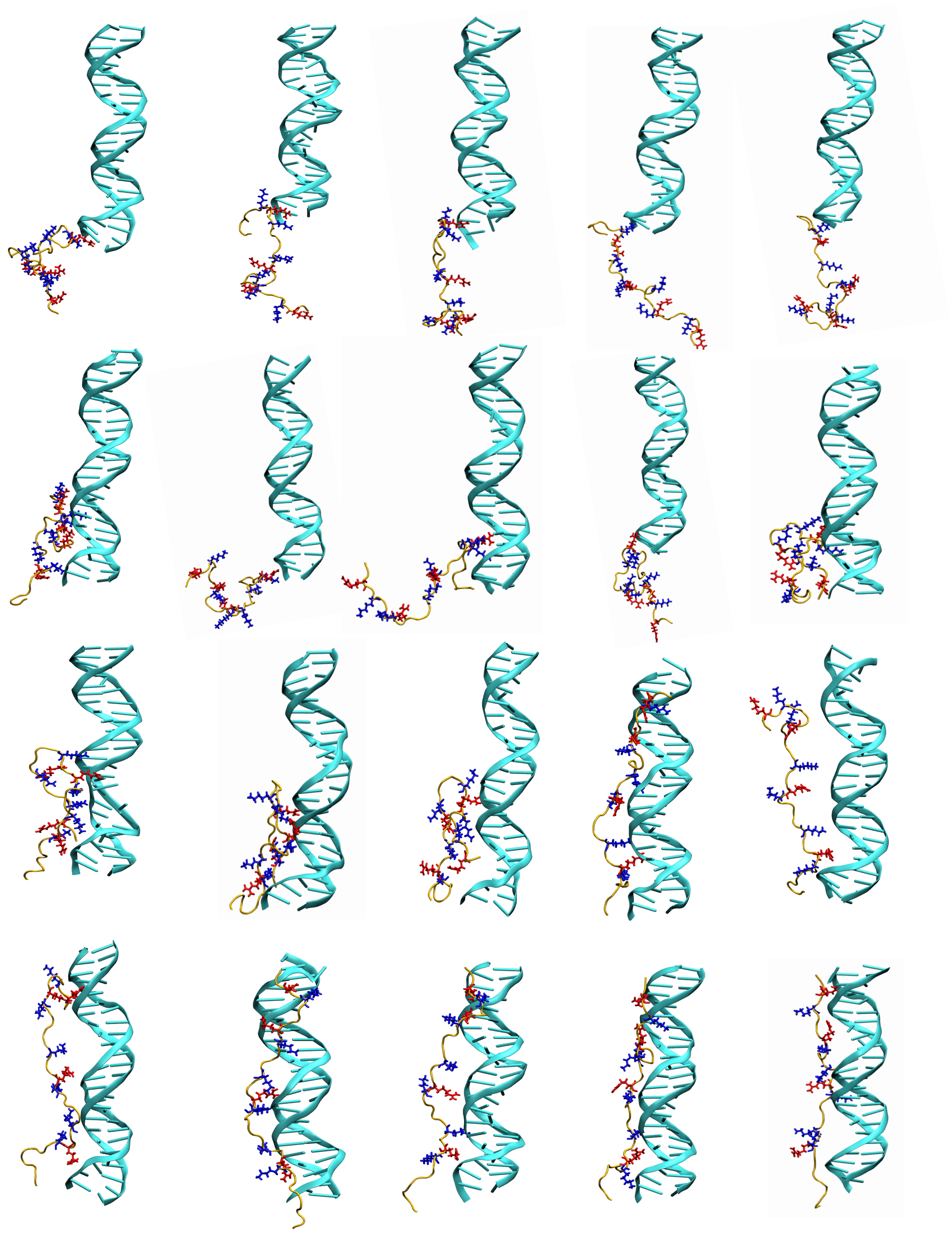

Figure S5: Alignment of long arginine and lysine amino acids with DNA major grooves in the absence of RNA.

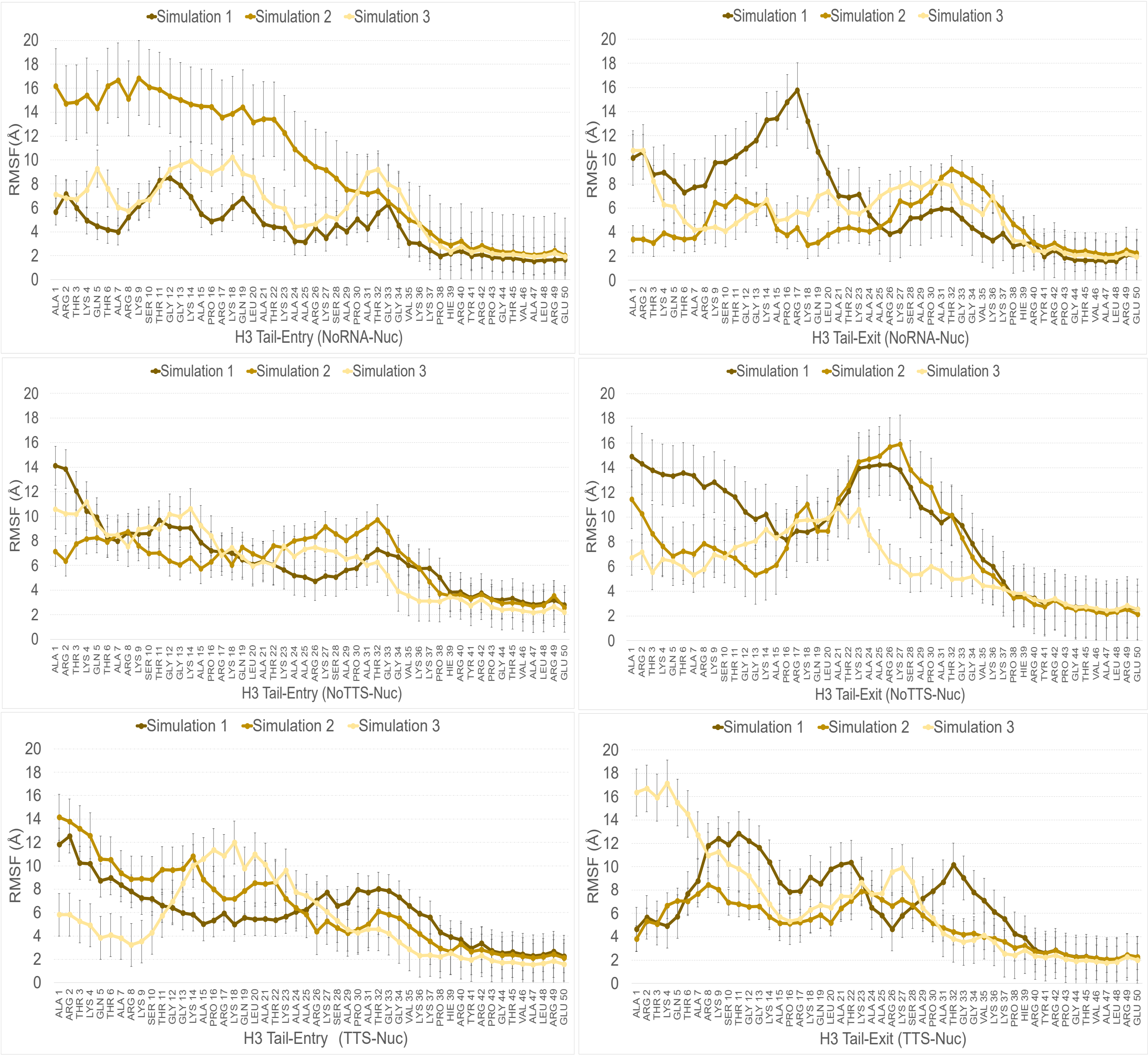


Figure S6: RMSFs of H3 tails in proximity of the entry/exit DNA for each simulation.

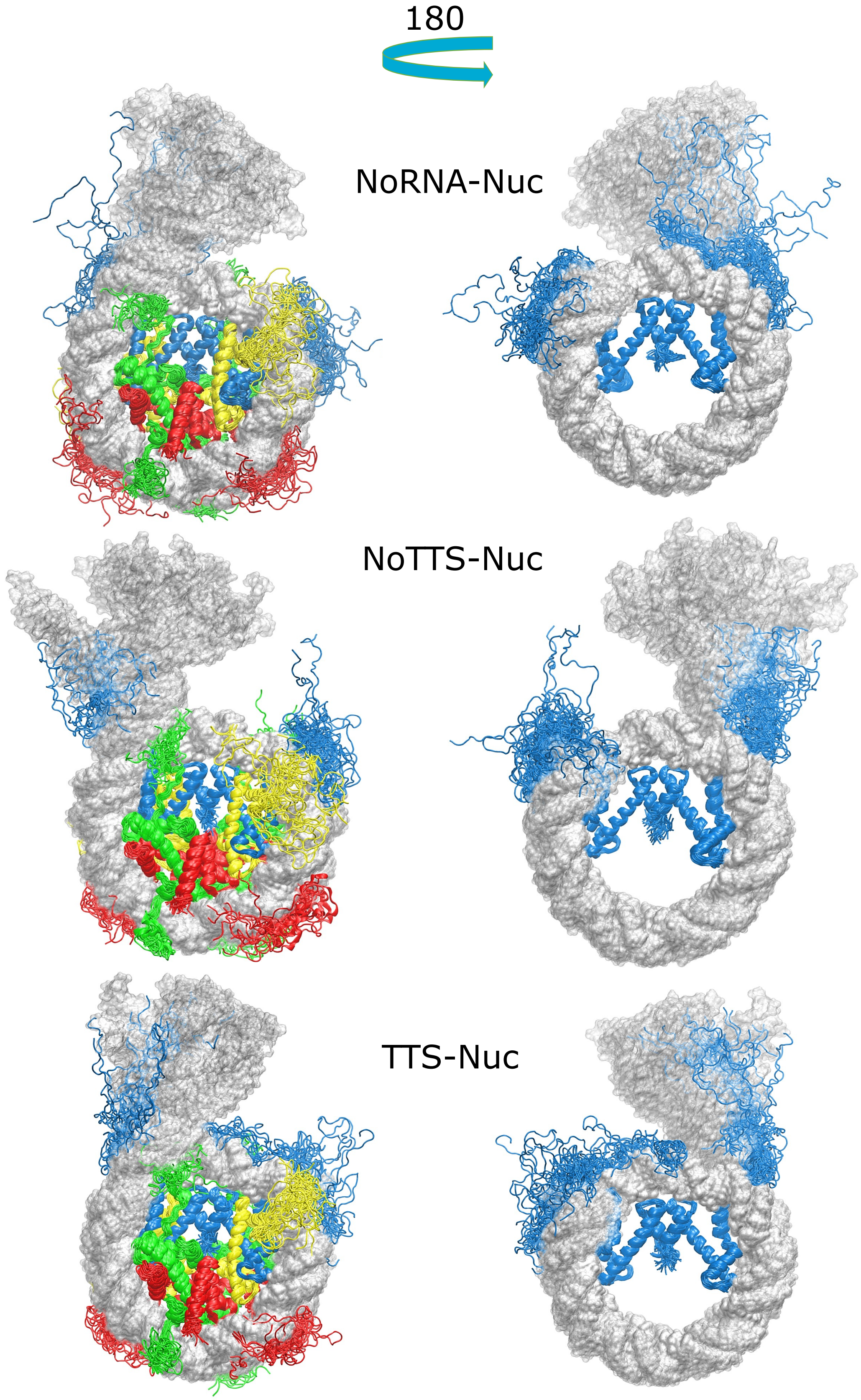


Figure S7: Dynamics of H3 tails interacting with DNA in NoRNA, TTS and NoTTS nucleosomes.

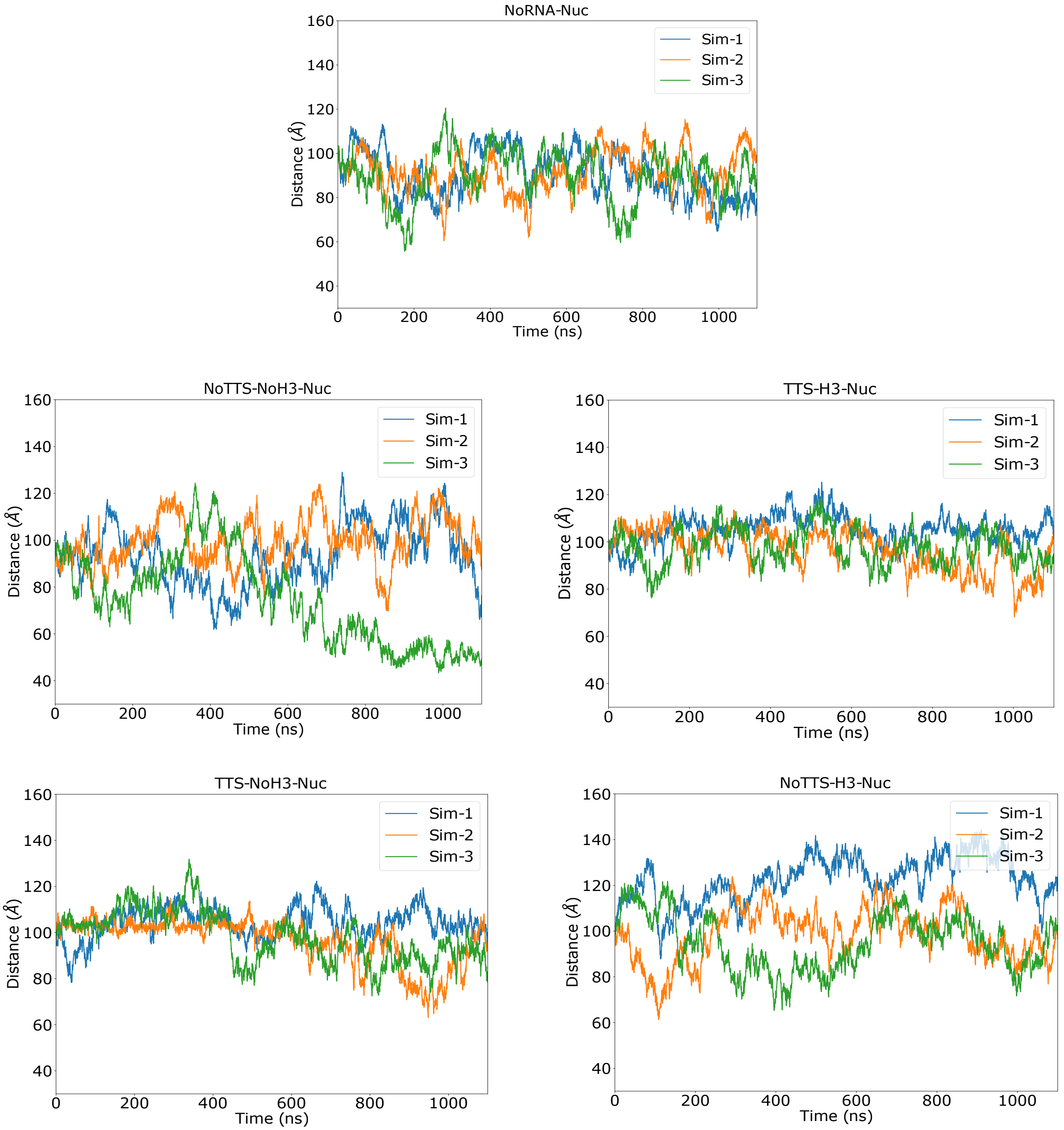


Figure S8: DNA end to end distance in individual simulations.

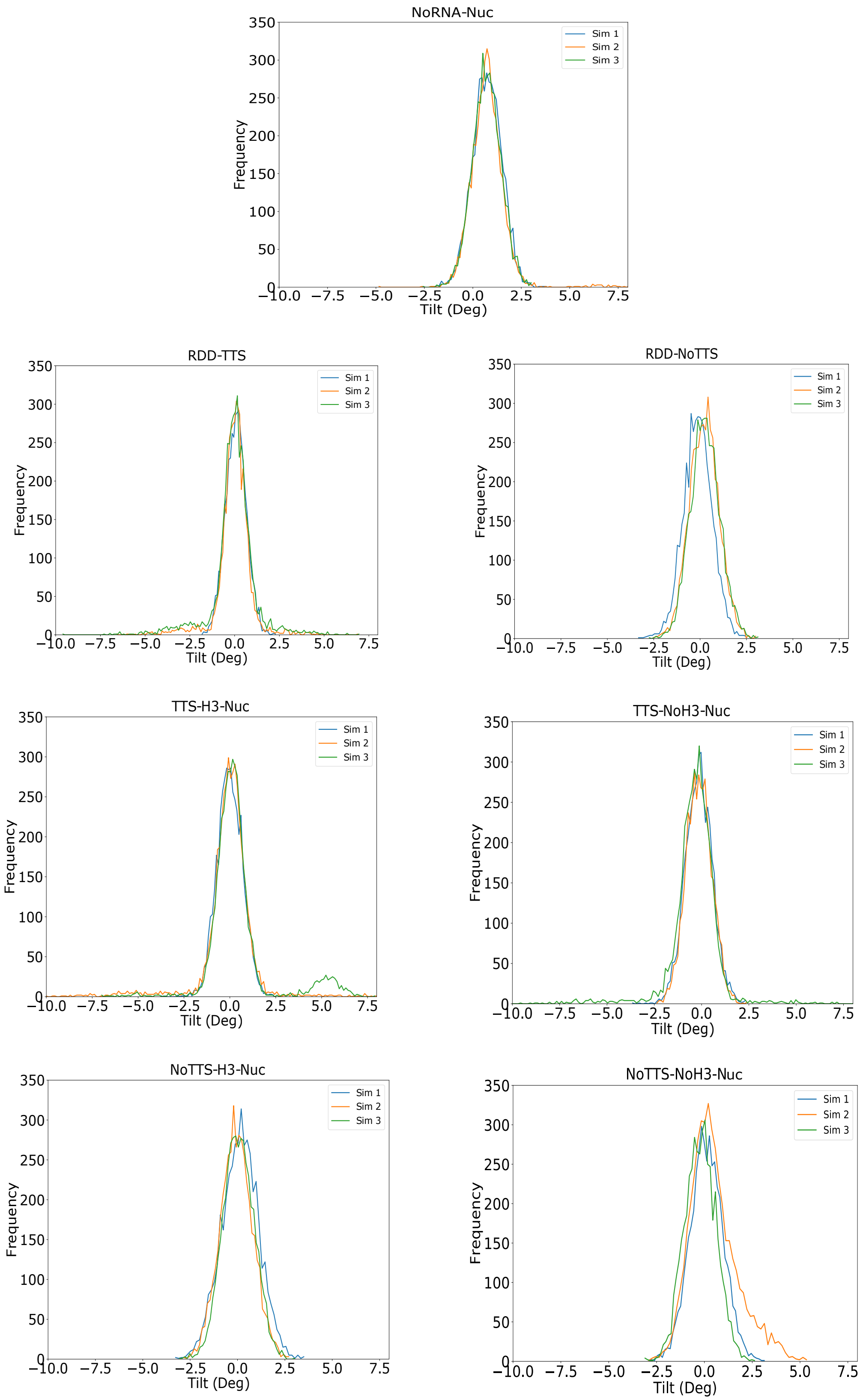


Figure S9: Distributions of the tilt parameter.

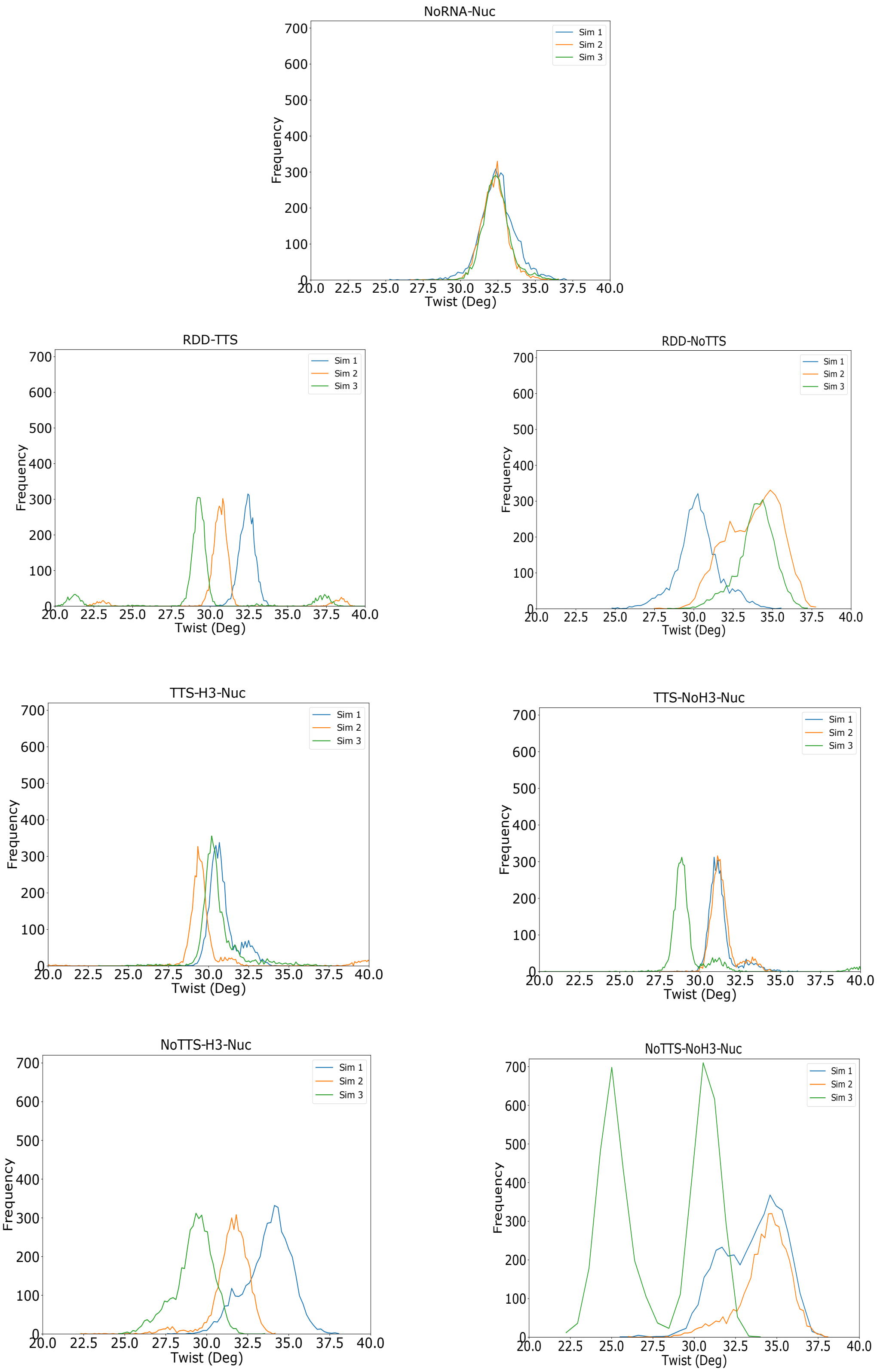


Figure S10: Distributions of the twist parameter.

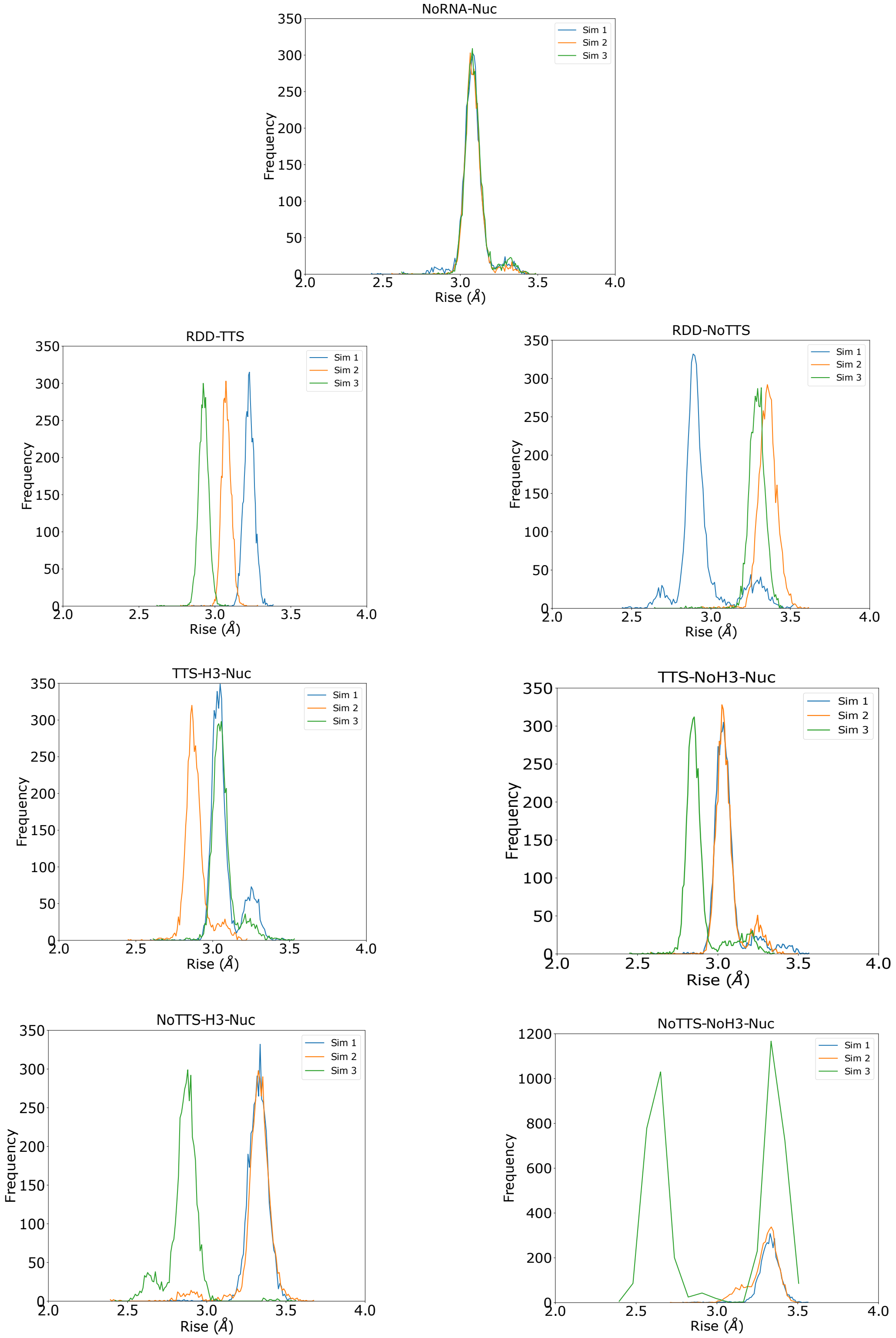


Figure S11: Distributions of the rise parameter.

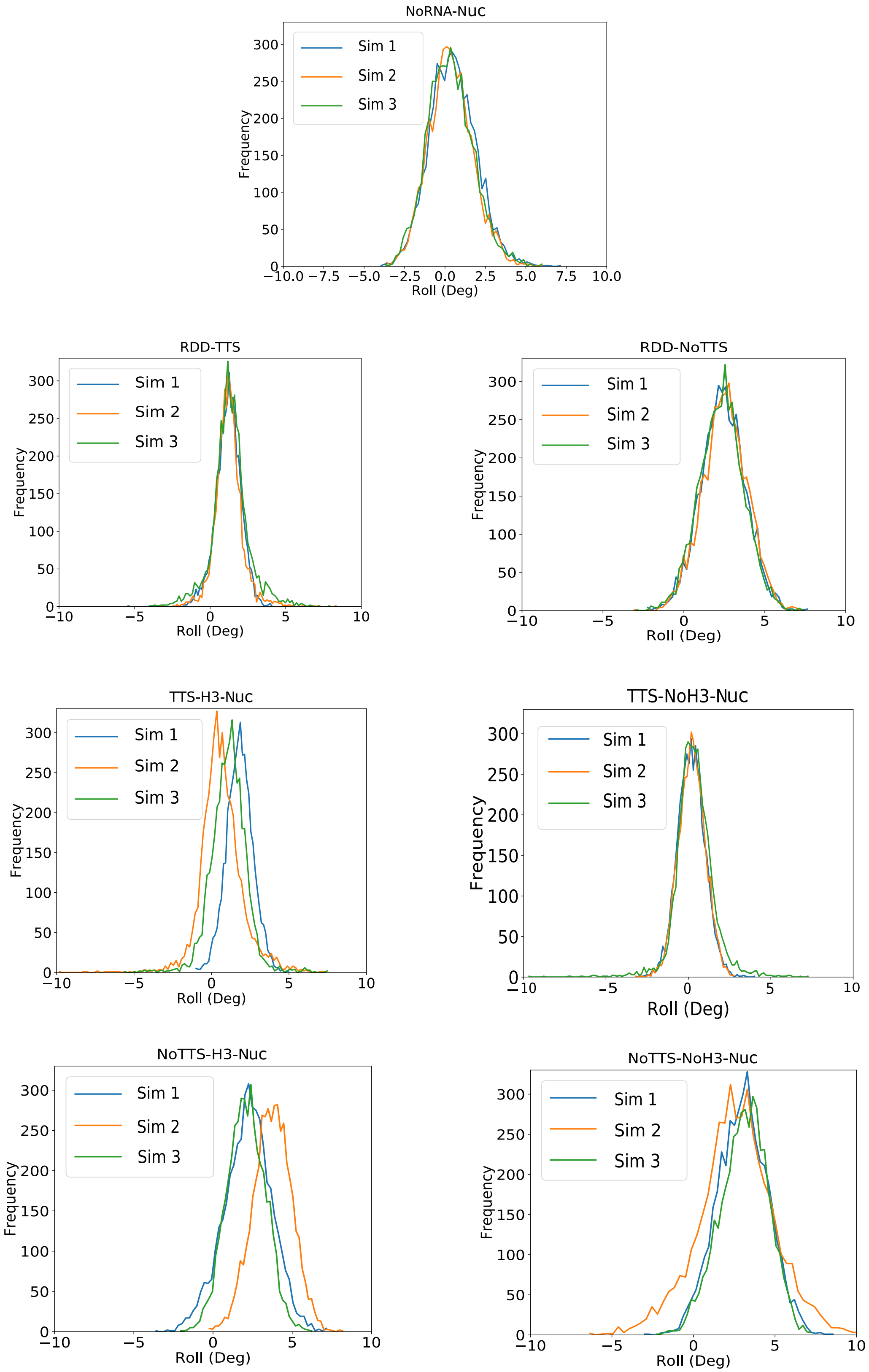


Figure S12: Distributions of the roll parameter.

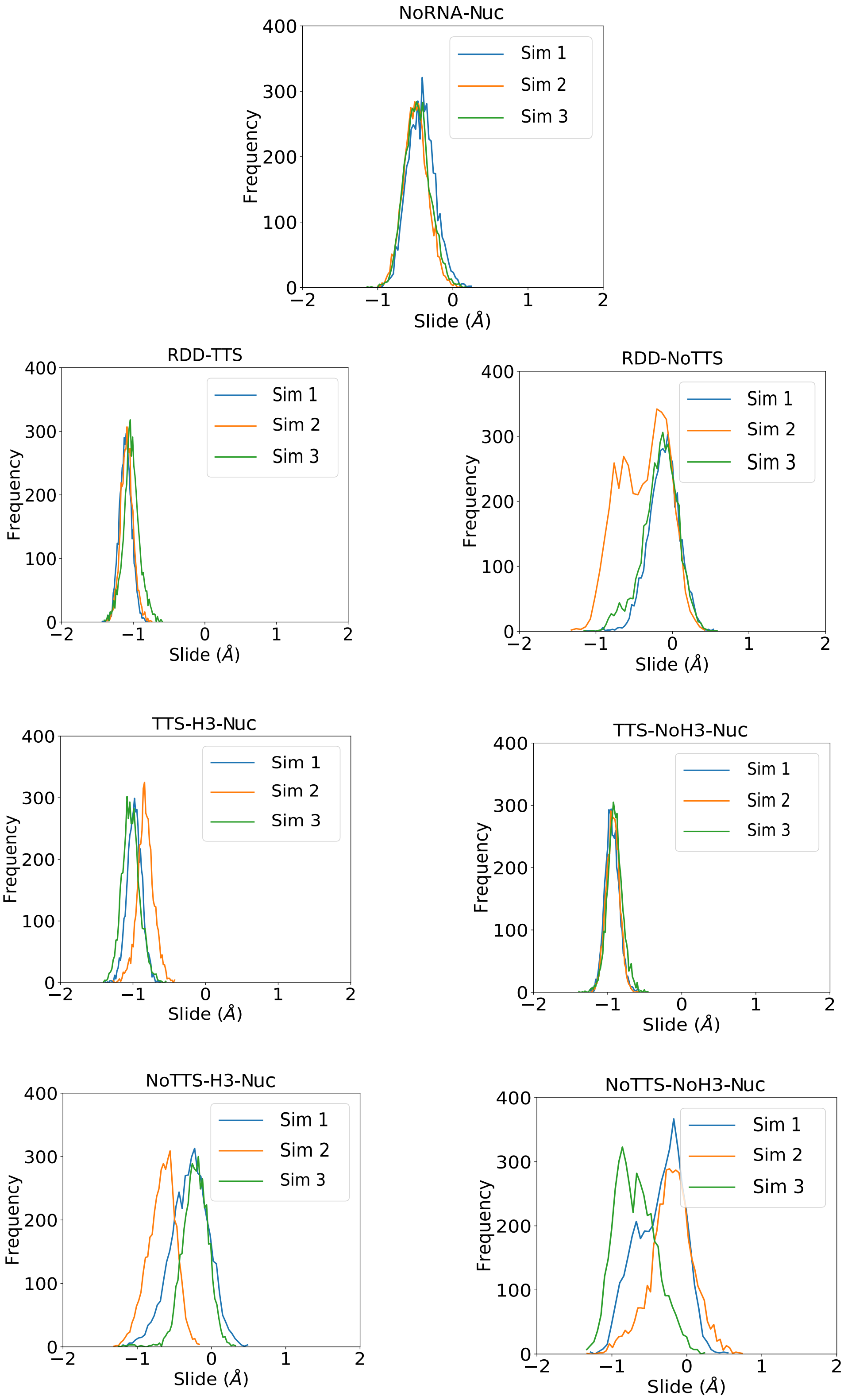


Figure S13: Distributions of the slide parameter.

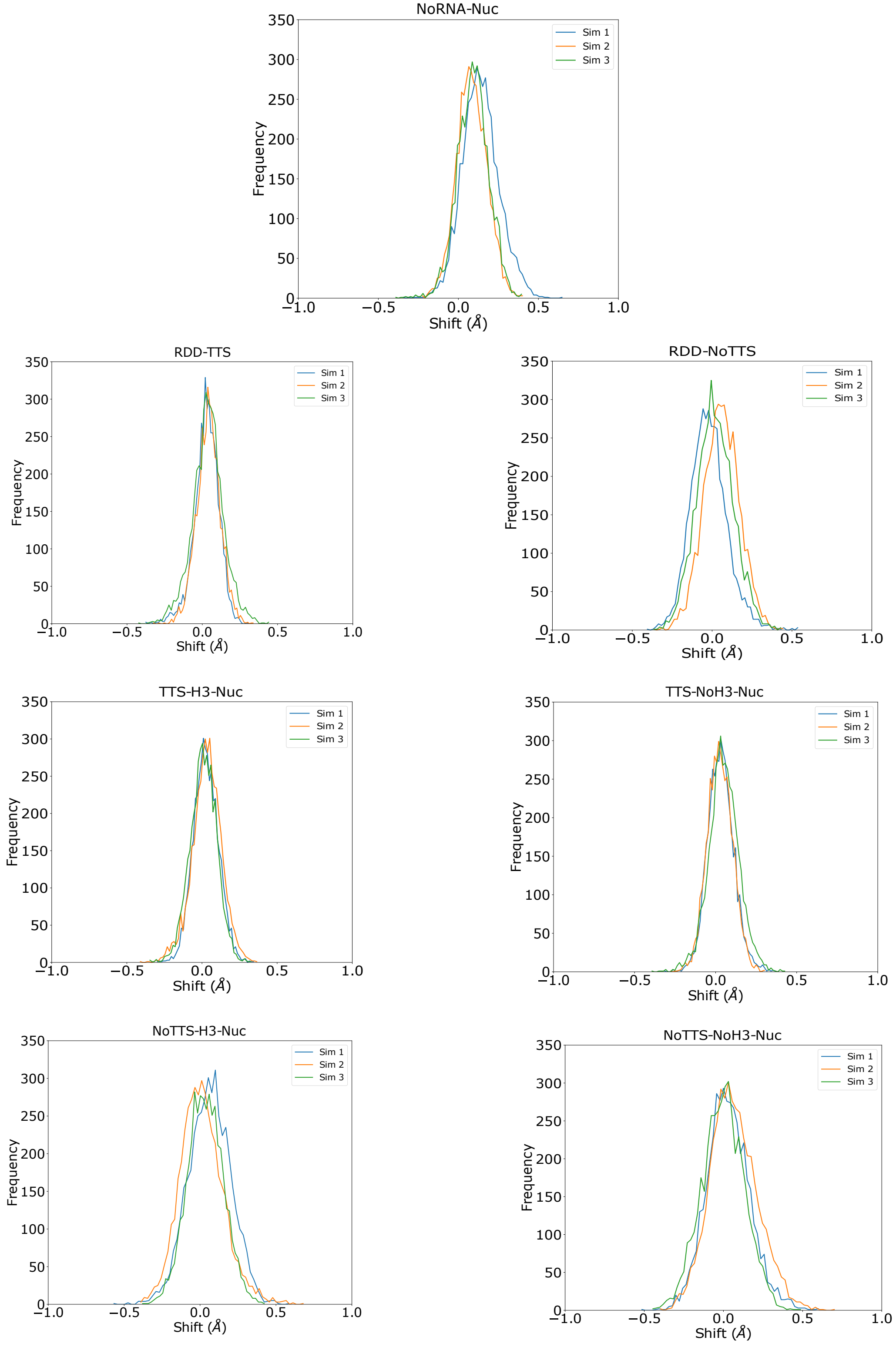


Figure S14: Distributions of the shift parameter.

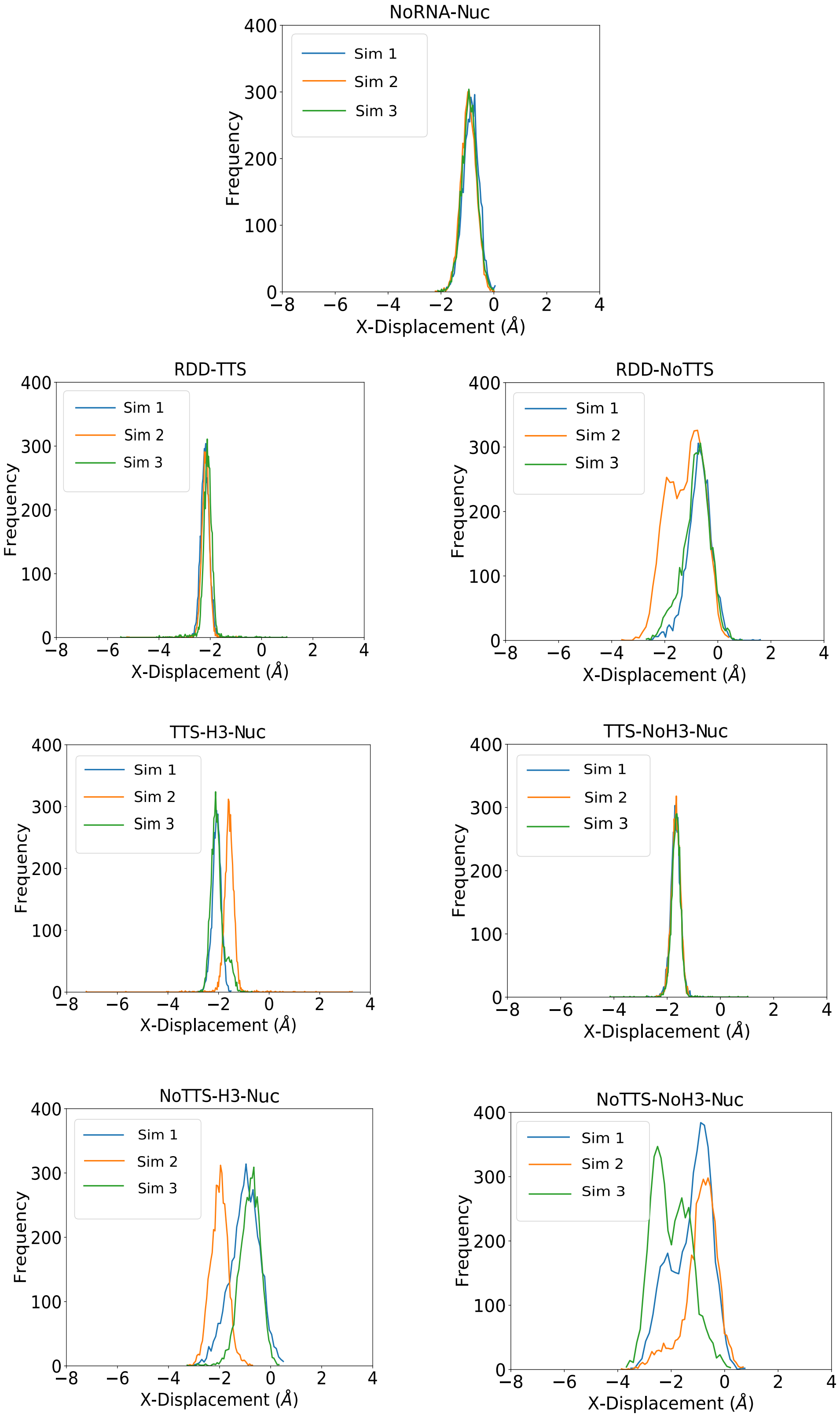


Figure S15: Distribution plots for helix X-displacement parameter.

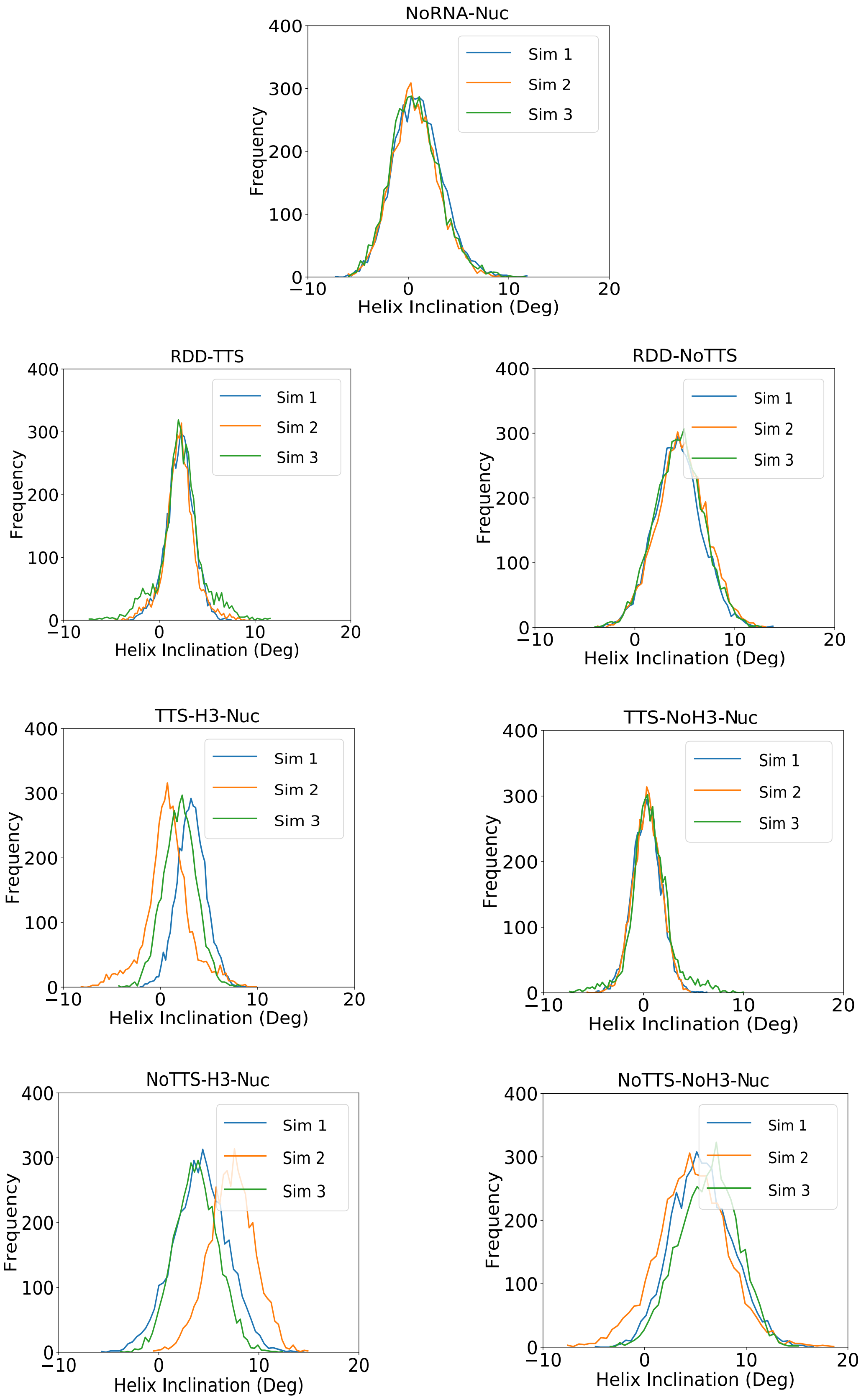


Figure S16: Distributions of the helix inclination parameter.

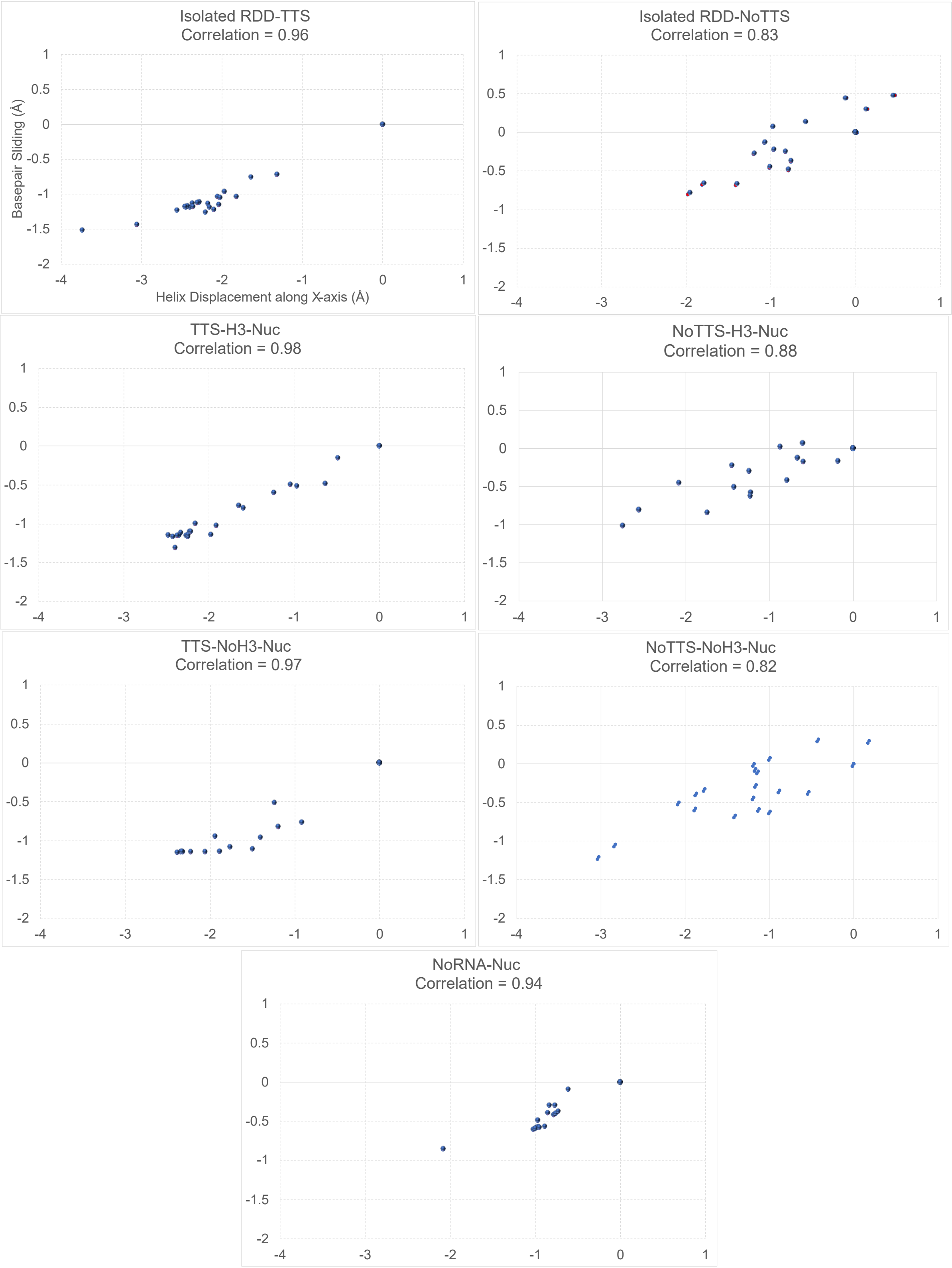


Figure S17: Helix X-displacement and base-pair slide are highly correlated.

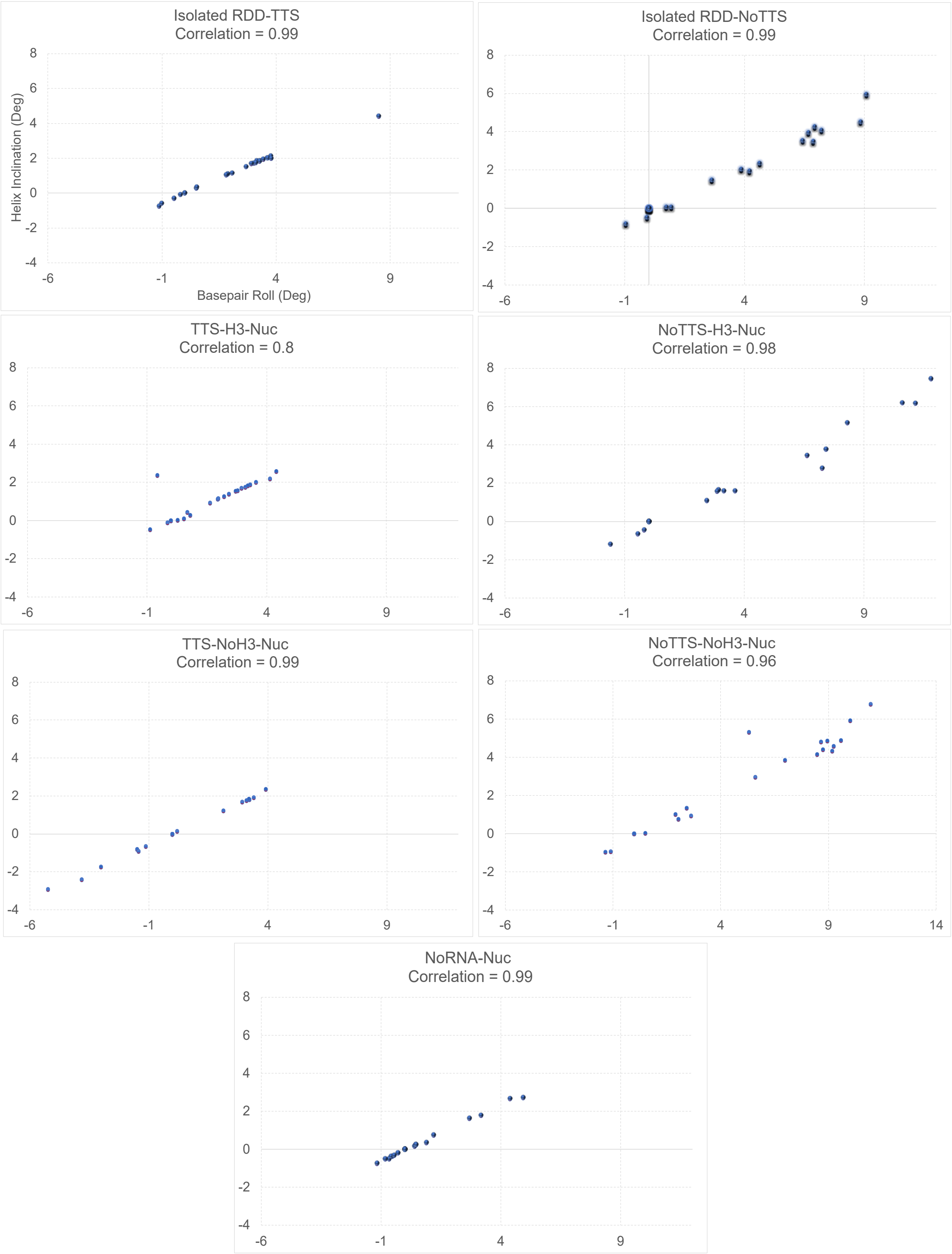


Figure S18: Helix inclination and base-pair roll are highly correlated.
